# Reduced anterior cingulate cortex volume induced by chronic stress correlates with increased behavioral emotionality and decreased synaptic puncta density

**DOI:** 10.1101/2020.08.31.275750

**Authors:** Keith A. Misquitta, Amy Miles, Thomas D. Prevot, Jaime K. Knoch, Corey Fee, Dwight F. Newton, Jacob Ellegood, Jason P. Lerch, Etienne Sibille, Yuliya S. Nikolova, Mounira Banasr

## Abstract

Clinical and preclinical studies report that chronic stress induces behavioral deficits as well as volumetric and synaptic alterations in corticolimbic brain regions including the anterior cingulate cortex (ACC), amygdala (AMY), nucleus accumbens (NAc) and hippocampus (HPC). Here, we aimed to investigate the volumetric changes associated with chronic restraint stress (CRS) and link these changes to the CRS-induced behavioral and synaptic deficits. We first confirmed that CRS increases behavioral emotionality, defined as collective scoring of anxiety- and anhedonia-like behaviors. We then demonstrated that CRS induced a reduction of total brain volume which negatively correlated with behavioral emotionality. Region-specific analysis identified that only the ACC showed significant decrease in volume following CRS (p<0.05). Reduced ACC correlated with increased behavioral emotionality (r=-0.56; p=0.0003). Although not significantly altered by CRS, AMY and NAc (but not the HPC) volumes were negatively correlated with behavioral emotionality. Finally, using structural covariance network analysis to assess shared volumetric variances between the corticolimbic brain regions and associated structures, we found a progressive decreased ACC degree and increased AMY degree following CRS. At the cellular level, reduced ACC volume correlated with decreased PSD95 (but not VGLUT1) puncta density (r=0.35, p<0.05), which also correlated with increased behavioral emotionality (r=-0.44, p<0.01), suggesting that altered synaptic strength is an underlying substrate of CRS volumetric and behavioral effects. Our results demonstrate that CRS effects on ACC volume and synaptic density are linked to behavioral emotionality and highlight key ACC structural and morphological alterations relevant to stress-related illnesses including mood and anxiety disorders.

**Highlights:**

1. Chronic restraint stress (CRS) decreases anterior cingulate cortex (ACC) volume
2. ACC and amygdala (AMY) volumes negatively correlate with behavioral emotionality
3. CRS decreased the strength and degree of the ACC structural covariance network
4. CRS increased the strength and degree of the AMY structural covariance network
5. PSD95 puncta density correlates with behavioral emotionality and ACC volume.

## 1 Introduction

Exposure to stressful environment triggers a variety of adaptive responses followed by the return to a homeostatic state. However, when stress exposure becomes chronic, the stress response gradually becomes maladaptive and results in detrimental effects (McEwen, 2017). Indeed, exposure to stressful situations can lead to emotional dysregulation (Hammen et al., 2009) and is a major risk factor in the development of psychiatric illnesses, including major depressive disorder (MDD) (Kessler, 1997) and anxiety disorders (Kessler et al., 2003). The mechanisms involved in the maladaptive response to chronic stress include functional dysregulation within the corticolimbic circuit (Hariri and Holmes, 2015). This circuit, which is highly conserved across species, encompasses the amygdala (AMY), hippocampus (HPC), nucleus accumbens (NAc) and regions of the prefrontal cortex (PFC) that include the anterior cingulate cortex (ACC). These four brain regions are key structures implicated in emotion regulation and functional dysregulations of and have all been implicated in MDD and anxiety disorders (Ai et al., 2015; Ge et al., 2019; Johnstone et al., 2007; Klauser et al., 2015; Rao et al., 2016; Rodríguez-Cano et al., 2014; Tang et al., 2013; Webb et al.; Ye et al., 2015; Zhao et al., 2014). This circuit is responsible for maintaining the homeostatic balance following stress exposure. The function of each brain region of the corticolimbic circuit is influenced by numerous efferent and afferent projections from other brain regions, collectively creating a network critical for a healthy stress response as well as in pathological deficits involved in stress-related illnesses, such as MDD (Kim et al., 2011; Kong et al., 2013).

Neuroimaging studies are valuable tools to identify macroscale structural and functional alterations associated with neuropsychiatric disorders. Numerous magnetic resonance imaging (MRI) studies have observed volumetric changes in corticolimbic brain regions as well as brainwide structural adaptations associated with MDD (McKinnon et al., 2009; Schmaal et al., 2017; Schmaal et al., 2020; Schmaal et al., 2016; Treadway et al., 2015). Particularly, MRI studies identified decreases in volume of cortical areas including the ACC (Hajek et al., 2008; Koolschijn et al., 2009b; Schmaal et al., 2017) and hippocampal regions in MDD patients (Campbell et al., 2004; Kaymak et al., 2010; Koolschijn et al., 2009a; Malykhin et al., 2010; McKinnon et al., 2009; Schmaal et al., 2016; Steffens et al., 2011; Videbech, 2004). Further studies have described progressive changes associated with cortical thinning and greater number of prior major depressive episodes (Macqueen et al., 2003; Treadway et al., 2015). However mixed results have been reported regarding structural alterations of the AMY and NAc. Increased, decreased or no changes in volume of the AMY have been described (Arnone et al., 2012; Frodl et al., 2003; Hamilton et al., 2008; Schmaal et al., 2016). Regarding the NAc, MRI structural findings are limited and inconsistent, where for example one study reported increased volume of this brain region associated with MDD (Abdallah et al., 2017), while another study has reported decreased NAc volume with lifetime MDD (Ancelin et al., 2019).

Postmortem studies attribute these volumetric changes identified using MRI to altered neuronal complexity and synaptic loss or gain in the PFC (Kang et al 2012) and the HPC (Boldrini et al., 2013; Cobb et al., 2016; Cobb et al., 2013; Stockmeier et al., 2004). More specifically, MDD is associated with decreases in size of neuron soma, dendritic architecture and synaptic loss in the PFC (Bianchi et al., 2005; Kang et al., 2012; Rajkowska et al., 1999). Gene and protein expression of synaptic markers were reported reduced in the PFC primarily focusing on the ACC (Scifo et al., 2018; Shukla et al., 2020) in the HPC (Duric et al., 2013), while increased in the NAc and the AMY (Labonté et al., 2017).

To better understand the pathophysiology of chronic stress-related disorders such as MDD, chronic stress-based animal models are used to investigate the biological mechanisms that may underlie changes resulting in the dysfunction of the corticolimbic system. Such models have been extensively investigated to assess depressive-like behaviours, (Banasr and Duman, 2007; Nikolova et al., 2018; Willner, 2017) antidepressant efficacy (Banasr and Duman, 2007; Prevot et al., 2019b) and to determine sex-specific pathophysiological mechanisms (Guilloux et al., 2011; Seney and Sibille, 2014). Synaptic and morphological changes are found in chronic stress models, e.g., dendritic atrophy and spine loss in regions of the PFC (Cook and Wellman, 2004; Eiland et al., 2012) and hippocampus (Cook and Wellman, 2004; Eiland et al., 2012; Hill et al., 2011; Li et al., 2011; Radley et al., 2004; Radley et al., 2005; Son et al., 2012; Vyas et al., 2002; Wellman, 2001) or hypertrophy in the AMY (Eiland et al., 2012; Hill et al., 2013; Vyas et al., 2006; Vyas et al., 2002). Most of these studies used the chronic restraint stress (CRS), which is the model we chose for the present study to establish links between synaptic, volumetric and behavioral alterations associated with CRS.

Recent advancements in rodent structural MRI have proven to be valuable in studying the morphological changes associated with chronic stress across experimental models (Anacker et al., 2016; Kassem et al., 2013; Lee et al., 2009; Magalhães et al., 2018; Nikolova et al., 2018). These studies allow for the simultaneous investigation of morphological changes in multiple brain regions and can thus facilitate the identification of whole-brain network adaptations to chronic stress. In support of this notion, we recently used MRI to link unpredictable chronic mild stress with volumetric and structural covariance network changes centered on the AMY with depressive-like behavior and altered synaptic marker density (Nikolova et al., 2018).

In the present study, we aimed to examine the effects of chronic stress at the structural level using MRI-measured volumes, focusing on key brain regions of corticolimbic circuit, the PFC, HPC, AMY and NAc. The role of these volumetric changes on behavior was investigated by identifying potential relationships between the chronic stress-induced volumetric changes in these brain regions and behavior (anxiety-, anhedonia-like and cognitive behavioral deficits). We then analyzed the consequences of chronic stress on structural covariance network patterns of these regions and sought to determine the underlying substrate of these changes by examining if regions with significant volumetric changes also showed altered synaptic density using immunohistochemistry. Finally, an unbiased exploratory whole brain volume analysis was performed in all animals to investigate the brain-wide structural changes associated with chronic stress exposure.

## 2 Methods

### 2.1 Animals

Eight-week old C57BL/6 mice (Jackson Laboratory, Bar Harbor, Maine, USA) were single housed under normal conditions with a 12hr light/dark cycle and provided with *ad libitum* access to food and water. All animals were habituated to the facility prior for 1 week to the start of the experimental procedures. All procedures were performed in accordance with institutional and Canadian Council on Animal Care (CCAC) procedural and ethical guidelines.

### 2.2 CRS

Animals were placed for a 1-hour period inside 50mL Falcon^®^ tubes with air holes cut into each end, twice daily (minimum 2hrs apart) for 2 or 5 weeks (n=12 per group). Control animals were handled daily (Prevot et al., 2019a; Prevot et al., 2019b). We opted for examining the effects of 2 and 5 weeks of CRS since these two time points are, on average, the shortest and longest most studied durations of CRS used the literature for determining the effects of chronic stress on synaptic and behavioral changes (Prevot et al., 2019b; Qiao et al., 2016). The 3 groups (50% females) were tested in all behavioral tests by an experimenter blinded to animal’s group assignment. In this study, physical changes and anhedonia-like behavior were assessed using classical parameters and tests used in chronic stress studies, such as the weight of the animals, the state of their coat and sucrose preference (Prevot et al., 2019b; Willner, 2017). We also assessed anxiety-like behaviors using the novelty suppressed feeding (NSF) and phenotyper tests since chronic stress induces consistent deficits in these tests across strains, sex and experiments as compared to less reliable anxiety tests such as in the elevated plus maze or open field tests (Prevot et al., 2019b).

### 2.3 Coat State Quality and Weight Measurements

Seven anatomical regions (head, neck, dorsal/ventral coat, forepaws, hindpaw and tail) on each mouse were assessed where each region was given a score of either 0, 0.5 or 1 from maintained (0) to unkempt (1). Values for each region were summed up into a single score for each animal. Weight was recorded at the same time as coat state quality to determine weight gain or loss (measured in grams) associated the CRS exposure (Nollet et al., 2013). Coat state assessment was performed weekly on day 7, 15, 21, 28, 35 (**Figure 1A**).

**Figure 1:**
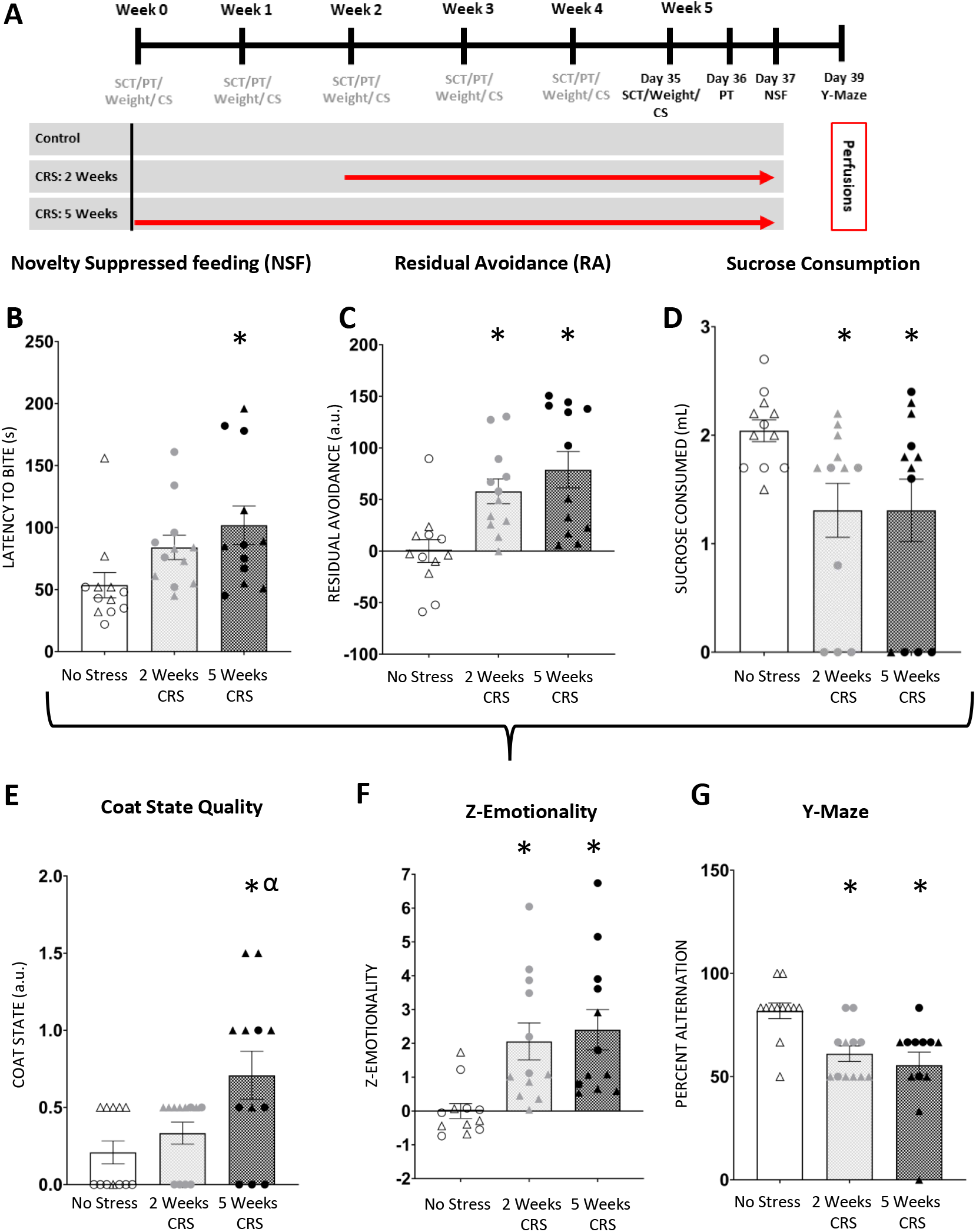
Chronic restraint stress exposure induces deficits in behavioral emotionality and working memory. (A) Schematic representation of the experimental design. Control animals and mice subjected to 2 or 5 weeks of chronic restraint stress (CRS) (n=12/group; 50% males) were tested in series of behavioral tests throughout the experiment (gray) and during the 5^th^ week of the experiment (black). CRS-exposed mice showed increased latency to bite in the (B) novelty suppressed feeding test (NSF) and increased residual avoidance in the (C) PhenoTyper test (PT). CRS animals also showed reduced sucrose consumption in the (D) sucrose consumption test (SC). While coat state degradation (E) was found significantly elevated in CRS animals. Overall CRS induced an increase in z-emotionality (F) calculated from the mean z-scores of the novelty suppressed feeding, PhenoTyper and sucrose consumption tests. CRS mice also displayed deficits in the (G) Y-maze working memory test. *p<0.05 compared to controls and ^α^p<0.05 compared to 2 weeks CRS. Individual males (Δ) and females (O) are represented in each figure.

### 2.4 Sucrose Consumption Test (SCT)

Before the start of CRS, animals were habituated to a 1% sucrose solution (Sigma, MO, USA) for 48 hrs and are then fluid deprived for 16 hours overnight. Sucrose consumption was measured the next morning for a 1-hour period. The same procedure was performed for water consumption after two days. For each follow-up SCT, animals were re-habituated to the solution (sucrose or water) for 24 hours, fluids were then removed for 16 hours and consumption was measured for a 1-hour test (Duric et al., 2017). This test was performed weekly on day 7, 15, 21, 28, 35 (**Figure 1A**).

### 2.5 PhenoTyper *Test (PT)*

The Noldus^™^ PhenoTyper (Leesburg VA, USA) is an observational home cage-like apparatus able to video track mice for extended periods of time (EthoVision 10 software). In the arena, there are two designated zones (food [6.5×15 cm] and shelter zones [10×10cm]) where amount of time spent was recorded during the animal’s dark cycle (19:00-07:00). In addition, an anxiogenic spotlight challenge appearing over the food zone begins 4 hours into the dark cycle for a 1-hour period (23:00-24:00). Testing in the PhenoTyper box was performed weekly on days 0, 7, 14, 21, 28, 35 of the CRS exposure. From previous studies we identified that stress-exposed and control animals spend equivalent amounts of time avoiding the food zone in favor of the shelter during the spotlight challenge. However, animals subjected to chronic stress continue to avoid the food zone after the challenge (Prevot et al., 2019b). This behavior, which we defined as “residual avoidance” (RA), is highly specific of chronic stress exposure (Prevot et al., 2019b). RA in the shelter zone was measured as the difference between the time spent during the light challenge and the sum of the time spent avoiding the lit zone (5 hours post-challenge). Formula for RA for each mouse as followed:

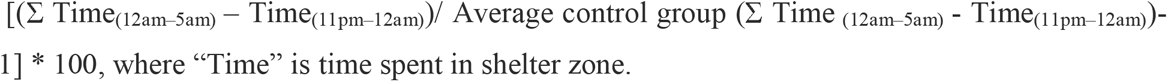

The RA measure provides a percentage of avoidance the animal exhibits after the white light challenge normalized to the control group. A positive RA value indicates the animal’s avoidance of the lit zone in favor of the shelter zone (Prevot et al., 2019b). This test was performed weekly the day after the sucrose preference test (**Figure 1A**).

### 2.6 Novelty Suppressed Feeding (NSF)

Mice are food deprived for 16 hours prior to testing. During testing, mice are placed into a novel arena (45 x 30 x 27 cm) with a single food pellet in the center under dim lit conditions (lux 28-30). The latency to approach and feed on the pellet is measured (in seconds). Similar test is performed in the animal’s home cage; home cage latency to feed was also measured to exclude experimental bias associated with appetite drive. This test was performed once at the end of the experiment on day 37 (**Figure 1A**).

### 2.7 Y-maze

The Y-maze is an apparatus with 3 arms (26 x 8 x 13cm, 120° apart) and sliding doors. Mice are habituated to the apparatus and allowed to explore the maze containing distal cues for 10 mins for 2 consecutive days. Mice are then trained to alternate for 7 successive trials and each trial is separated by an inter-trial interval (ITI) period of 30s. Each trial begins with a 30s period in the “start-box” after which the door is slid open allowing the animal to explore one side of the arm. The door to the arm the animals entered is immediately closed for a 30s period and choice of arm is recorded. Mice return to the “start-box” and similar consecutive trials proceed. The testing phase follows the same procedure as training phase, except that an ITI of 90s is used, and with the addition of a 8^th^ trial with a 5s ITI to assess for potential motivational loss. In the event of a lack of alternation on that trial, animals are removed from the analysis. All animals alternated in the last trial in this study. Data is calculated as the percent mean alternation rate (number of alternation/number of trials × 100). This is Y-maze spontaneous alternation task where the animals use spatial cues. No food reward was given. CRS animals were previously shown to display deficits in this working memory task (Prevot et al., 2019a). This test was performed once at the end of the experiment on day 39 (**Figure 1A**).

### 2.8 Mouse Brain Tissue Preparation

At the end of the experiment, mice were anesthetized with avertine (125mg/kg, i.p.) and intracardially perfused using a Pharmacia minipump at a rate of approximately 100mL/hr with 30 ml of 0.1M PBS containing 10U/mL heparin and 2mM ProHance (Bracco Diagnostics, NJ, USA), followed by 30 ml of 4% paraformaldehyde solution containing 2mM ProHance (Cahill et al., 2012; Lerch et al., 2011). After trans-cardiac perfusion, mouse bodies were decapitated. Skin, muscles, lower jaw, ears, and the cartilaginous nose tip were removed. The skull containing the brain was placed in a post-fixative solution of 4% PFA + 2mM ProHance overnight at 4°C. Samples were then transferred into a buffer storage solution of 0.1M PBS with 2mM ProHance and 0.02% sodium azide for a minimum of one month (De Guzman et al., 2016) using a 7.0 Tesla MRI scanner (Agilent Inc., Palo Alto, CA) as in Anacker et al. (2016).

### 2.9 MRI Acquisition and Preprocessing

Brain preparations and scanning was performed as described in Nikolova et al (2018). However, regional volume was parcellated into 182 brain regions excluding ventricles (instead of 159 as in Wheeler et al. (2015) and adapted from previous rodent MRI studies (Dorr et al., 2008; Richards et al., 2011; Steadman et al., 2014; Ullmann et al., 2013). To minimize multiple testing burden, all our MRI analyses focused on region of interest (ROI) volumes averaged across hemispheres. Analyses were conducted using deformation-based morphometry to determine the absolute brain volumes (in mm^3^). First, we conducted an initial *a priori* analysis focusing on the effects of chronic stress exposure on the volumes of 4 corticolimbic brain regions. This required summation of multiple subregions for the ACC (cingulate Area 24a, a’, b, b’), and hippocampal formation (CA1, CA2, CA3 and dentate gyrus). None was necessary for the NAc and AMY. Second, we tested a follow-up *a priori* analysis focusing on 26 MDD associated brain regions as in (Nikolova et al., 2018). For consistency, all these adjusted brain regions were used in the first *a priori* analysis, Pearson correlations and structural covariance analysis. Third, we conducted a whole-brain unbiased analysis in which these regions were separated into 182 regions of interest (ROIs).

### 2.10 Structural Covariance

Using the 182 ROIs, correlational matrices were created separately for each group. Based on previous studies, all negative correlations were removed (Nikolova et al., 2018). All matrices were thresholded over a range of density thresholds (0.05-0.4) that identified the top 5-40% strongest connections by increments of 1%. All thresholded matrices were converted to weighted graphs. We tested pairwise group differences (control vs. CRS 2 weeks, CRS 2 weeks vs. CRS 5 weeks, control vs. CRS 5 weeks) in regional degree and degree rank across density thresholds from 0.05 to 0.40 in increments of 0.01. Significance was determined using permutation testing (n=10,000) and defined as ≥ 5 consecutive density thresholds with p ≤ 0.05. Analyses were performed for the ACC, HPC, NAc and AMY, and regional volumes were adjusted for sex and total brain volume (refer to Nikolova et al. (2018)). Cytoscape (v3.7.1, Systems Biology, Seattle, USA) was used to visualize these networks at the lowest density threshold where observed between-group effects were significant. Subnetwork visualization of ROIs with significant structural covariance network changes (ACC and AMY) was created using the Edge-Weighted Spring Embedded Layout option in Cytoscape, as in Kamada and Kawai (1989), where each node represents the volume of a particular ROI and each edge represents the strength of correlation in volume between interconnected nodes.

### 2.11 Fluorescence Immunohistochemistry

After *ex-vivo* MRI, brains were extracted from the skull and placed in a 30% sucrose solution for cryoprotection. Free-floating sections (30μm thickness) were cut using a cryostat (Leica, Wezlar, German) and placed in a cryoprotectant solution (sucrose 30%, polyvinyl-pyrrolidone-40 (1%), 0.1M phosphate buffer (PB) and ethylene glycol 30%) for storage at −20°C until immunohistochemistry. Three free floating sections from the ACC (coordinates from Bregma 1.41 to 1.53mm) or the basolateral amygdala (BLA) (coordinates from Bregma −1.31 and −1.43 mm) were used for the immunohistochemistry. The sections were selected to represent anterioposterior coordinates as similar as possible for each animal. Sections were rinsed in 1X PBS phosphate-buffered saline (PBS, 20min) at 4°C and incubated in a 0.01M sodium citrate in distilled H_2_O at 80°C for 15 minutes. Solution was allowed to cool to room temperature and sections placed in a 0.3% X-100-Triton solution for 20 minutes followed by 1 hour in 20% donkey serum in PBS. Sections were incubated overnight at 4°C in PBS containing 2% donkey serum and primary antibodies for post-synaptic density protein 95 (PSD95; rabbit host, 1:100, Cell Signaling Technology, Danvers, MA, product #2507, lot 2), and vesicular glutamate transporter 1 (VGLUT1; guinea pig host, 1:500, Synaptic Systems, Goettingen, Germany, product #135304, lot 135304/31). Sections were rinsed in PBS and incubated at 4°C in PBS containing 2% donkey serum and secondary antibodies conjugated to anti-rabbit Cy3 (Jackson ImmunoResearch Inc., West Grove, PA) and anti-guinea pig CF405M (Biotium, Hayward, CA), which were used to detect PSD95, and VGLUT1 immunoreactivity, respectively (2 hours, donkey host, 1:500 for all). Sections were placed through a final wash in PBS for 20 mins. Sections were mounted onto Superfrost plus gold slides (Fisher Scientific, Massachusetts, United States), coverslipped (Vectashield Antifade Mounting Media, Vector Laboratories, Burlingame, CA), and stored at 4°C until imaging and puncta quantification analysis

### 2.12 Confocal Microscopy and Puncta Analysis

Confocal imaging of the ACC and BLA using an Olympus IX83 inverted microscope equipped with a spinning disk confocal unit and Hamamatsu Orca-Flash4.0 V2 digital CMOS camera using a 60X 1.4 NA SC oil immersion objective. SlideBook 6 (Intelligent Imaging Innovations, Denver, CO) was used to capture images. All images used for quantification were obtained from acquisition of approximately 40 stacks (covering ~4μm thickness) of top layer imaged. Image z-stacks were taken with a step size of 0.1 μm based on Nyquist rate to acquire the minimal optimal sampling density. Using stereological available tools in SlideBook 6, five image stacks (50×50×4 μm) were randomly taken for each hemisphere of the ACC (focusing on cingulate area 24) and the amygdala (focusing on the BLA). All images were taken using the same exposure settings for each channel and blind deconvolution was performed using Autoquant (Media Cybernetics, Inc., Rockville, MD). Puncta analysis was conducted using IMARIS v9.5.1 software (BitPlane, Badenerstrasse, Zurich, Switzerland) which compiles z-stacked images into a 3D rendering. Quality parameters, a threshold setting for the intensity of light at the center of each spot, was set to for optimal puncta density quantification for PSD95 (250 for ACC and 150 BLA) and for VGLUT1 (200 for ACC and 650 for BLA). A background subtraction with a maximum pixel diameter (xy-axis) of 0.35 and a point spread function width (z-axis) 0.5 was set. All results were calculated as the density in μm^3^.

### 2.13 Statistical analyses

Using SPSS statistical software (IBM, NY), analysis of variance (ANOVA), followed by Fisher’s post-hoc analysis, was performed on all behavioral tests. Repeated measures ANOVA was performed on the PhenoTyper, sucrose consumption, weight and coat state quality longitudinal assessments. A z-score normalization was performed to reduce behavioral data into one score for emotionality behavior. This calculation identifies the standard deviation of each group with respect to the mean of the controls. Z-scores were calculated from data obtained on the last testing in the PhenoTyper (residual avoidance), sucrose consumption, and NSF latency to bite tests and were averaged to create a z-emotionality score (Guilloux et al., 2011). Our *a priori* hypotheses of stress-induced volume changes in the 4 corticolimbic brain regions were tested using SPSS and separate ANCOVAs, including ROI volume as the dependent variable and total brain volume (TBV) and sex as regressors, followed by Fisher’s *post hoc* tests. Pearson correlation was used to assess relationships between z-emotionality and absolute volume change for the amygdala, ACC, NAc and hippocampus (with/without adjusting for TBV and sex). Similar analyses were performed with the 26 MDD-associated brain regions and with all 182 brain regions, and results were corrected for multiple comparisons using false discovery rate (FDR, q<0.05). Associations between ROI volumes, adjusted for TBV, and Z-emotionality scores were tested in R using Pearson correlation followed by FDR correction (q<0.05). Similarly, associations between TBV-adjusted ROI volumes and percent alternations in the Y-maze test were assessed using a multi-trial binary logistic regression (p<0.05). Finally, Pearson correlation was used to assess links between changes in synaptic puncta of PSD95 and VGLUT1 and ROI volume and z-score emotionality in SPSS, while logistic regression was used to assess links between Y-maze percent alternation and puncta density in R.

## 3 Results

### 3.1 CRS increases z-emotionality and decreases Y-maze performances assessed at 2 and 5 weeks

**Supplementary Material** details results obtained with longitudinal assessments of behavioral changes in the PT (**Supplementary Figure 1)** as well as sucrose consumption, weight and coat state changes, performed weekly to monitor efficacy of CRS exposure throughout the experiment (**Supplementary Figure 2)**. Differences between sexes were analyzed for all the tests (**Supplementary Figures 3 and 4**).

During the final week of testing, endpoint tests included the coat state assessment, NSF, PT and sucrose consumption to assess the effects of CRS on behavioral emotionality. In the NSF test, ANOVA of latency to bite revealed a significant main effect of stress (F_(2, 30)_ = 4.149; p < 0.05; **Figure 1B**). *Post hoc* analysis identified a significant increase in latency to bite between the animal group subjected to 5 weeks of CRS compared with the control mouse group (p<0.05). No significant main effect of sex and stress x sex interaction were found (**Supplementary Figure 3**). Appetite drive as measured by home-cage latency to feed revealed no significant differences between groups.

In the PT, RA in the shelter zone was calculated as a measure of anxiety behavior as in Prevot et al. (2019). ANOVA of shelter RA revealed a significant main effect of stress (F_(2, 30)_ = 23.788; p < 0.0001), sex (F_(1, 30)_ = 37.526; p < 0.0001) and stress x sex (F_(2, 30)_ = 11.259; p < 0.001) explained by a significant increase in RA in mice exposed to both 2 and 5 weeks of CRS as compared to controls (p<0.05) (**Figure 1C**). We also identified a significant increase in RA in 2 and 5 weeks CRS exposed female mice as compared to female controls and in CRS male mice compared to male controls (**Supplementary Figure 3**). Further ANOVA revealed a greater RA in females compared to males in mice exposed to both 2 and 5 weeks of CRS (**Supplementary Figure 3**).

The sucrose consumption test was measured as an index of anhedonia-like behavior and was conducted weekly (**Supplementary Figure 2**). The last sucrose consumption test performed on week 5, revealed a significant main effect of stress (F_(2, 30)_ = 4.403; p < 0.05) and sex (F_(1, 30)_ = 7.005; p < 0.05) but no stress x sex interaction was found. *Post hoc* analysis identified a significant decrease in sucrose consumption in mice exposed to both 2 and 5 weeks of CRS compared to control mice (p<0.05; **Figure 1D)**. Furthermore, a significant decrease in sucrose consumption in females exposed to 2 and 5 weeks of CRS as compared with control females (p<0.05) was found. This was not evident in males (**Supplementary Figure 3**).

The NSF, PT and sucrose consumption tests were z-scored and compiled into a z-emotionality score. We found a significant main effect of stress (F_(2, 30)_ = 12.344; p < 0.0001), sex (F_(1, 30)_ = 17.604; p < 0.0001) and stress x sex interaction (F_(2, 30)_ =4.449; p<0.05). P*ost hoc* analysis identified a significant increase in z-emotionality score in mice exposed to 2 and 5 weeks of CRS compared to control mice (p<0.05; **Figure 1F**). Increased z-emotionality score was significant in female mice exposed to 2 and 5 weeks of CRS compared to males exposed or not to CRS (p<0.05; **Figure 2A**). Further, males exposed to 5 weeks of CRS revealed a significant increase in z-emotionality as compared to the male no stress group (p<0.05; **Figure 2A**).

**Figure 2:**
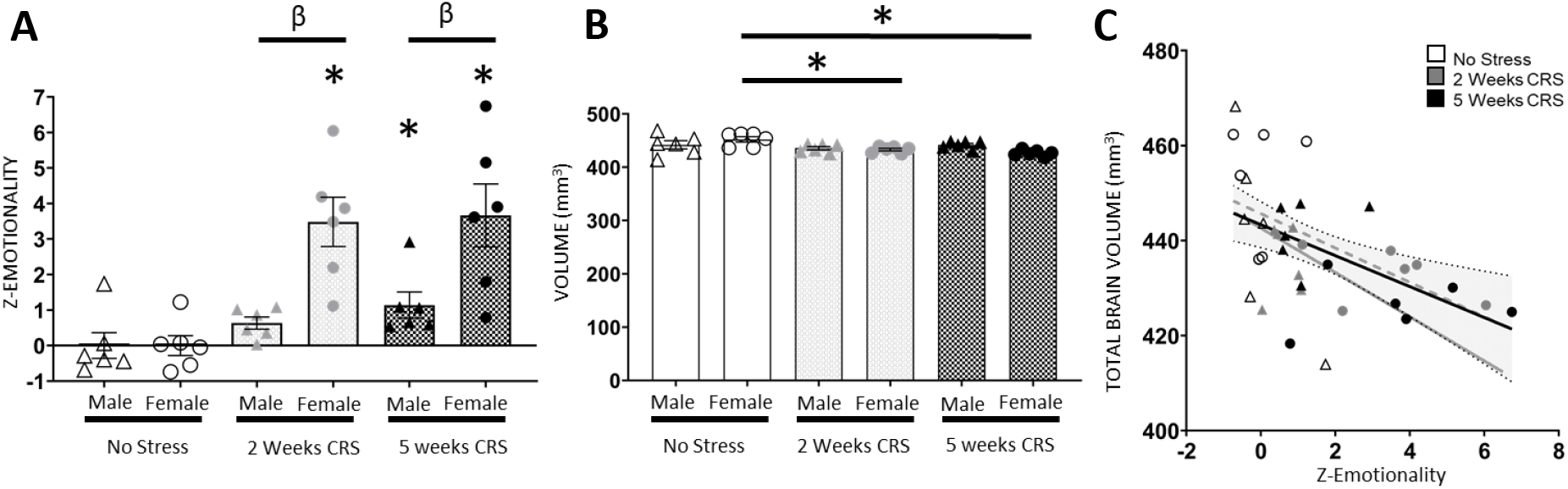
Chronic stress-induced deficits in behavioral emotionality and total brain volume changes are greater in female mice. Male and female mice were exposed or not to chronic restraint stress (CRS) (n=6/group). (A) Overall z-emotionality scores were calculated for each group and each sex. We found a decreased in (B) total brain volume (TBV) in female mice exposed to 2 and 5 weeks of CRS. Scatter plot showing the correlation (95% confidence interval) between (C) TBV changes and z-emotionality. A significant negative correlation was found when all animals (r=-.49; P=0.002; black line), or only females (r=-0.59; P=0.01; dotted grey line) were considered, but not when only males (grey line) considered. Individual males (Δ) and females (O) are represented in each figure. *p<0.05 as compared to respective controls; ^β^p<0.05 as compared to males.

Lastly the Y-maze test was used to assess effects of CRS on cognition, specifically working memory. We found a significant effect of stress (F_(2, 30)_ = 10.524; p < 0.001) and sex (F_(1, 30)_ = 5.909; p < 0.05) on percent alternations, but no stress x sex interaction. *Post hoc* analysis revealed a significant decrease in percent alternations in mice exposed to 2 or 5 weeks of CRS as compared to controls (p< 0.05; **Figure 1G**); this effect was significant only in males (**Supplementary Figure 3**). Analysis of coats state quality at 5 weeks showed a significant main effect of stress (F_(2, 33)_ = 5.797; p < 0.01). *Post hoc* analysis revealed a significant increase in coat state score between mice exposed to 5 weeks of CRS as compared to controls and 2 weeks CRS exposed mice (p<0.05; **Figure 1E)**.

### 3.2 Total brain volume decreases in CRS-exposed mice and correlates with z-emotionality

Total brain volume significantly decreased (F_(2,30)_ =5.666; p<0.001) in animals exposed to 2 or 5 weeks of CRS as compared to controls. There was no main effect of sex, however we found a stress x sex interaction (F_(2,30)_ =4.298; p<0.05) revealing that only females showed a significant decrease in TBV when exposed to 2 or 5 weeks of CRS (p<0.05, for both CRS group relative to control group, **Figure 2B**). Pearson’s correlation revealed a significant negative correlation between TBV and z-emotionality score across groups (r= −0.495; p=0.002, **Figure 2C).** While such correlation was detected in females (r= −0.608; p=0.007, **Figure 2C)**, no significant correlation was found in male mice (r= −0.336; p=0.173, **Figure 2C).** Using binary logistic regression, we found that TBV did not predict Y-maze performance (p>0.05).

### 3.3 ACC volume decreases in CRS-exposed mice and corticolimbic limbic regional volumes negatively correlate with z-emotionality

The initial *a priori* analysis using ANCOVA revealed significant PFC volume changes following CRS (F_(2,31)_ =3.826; p<0.05) using TBV as a covariate and sex as a cofactor; however, this effect was primarily driven to a significant effect of stress on the ACC volume (F_(2,31)_ =4.053; p<0.05; **Figure 3A**) as no other PFC regions showed significant changes. *Post hoc* analysis revealed a significant decrease in ACC volume in mice exposed to 2 or 5 weeks of CRS, compared to control mice (p<0.05). However, no significant changes in volume were found in the AMY, NAc and HPC following CRS (**Figure 3**). The absolute volumes of the ACC (*r* = −0.56; p=0.0003; **Figure 3B**), AMY (*r* = −0.35; p=0.03; **Figure 3F**) and NAc (*r* =-0.41; p= 0.01; **Figure 3H**) negatively correlated with z-emotionality score. The HPC volume did not correlate with z-emotionality score. This analysis was performed using absolute volume of each brain region and did not survive when TBV was included in the analysis as a covariate and sex as a cofactor (p>0.05). Using binary logistic regression, we found that volume of the ACC, AMY, NAc and HPC did not predict Y-maze performances **(Supplementary Table 3).**

**Figure 3:**
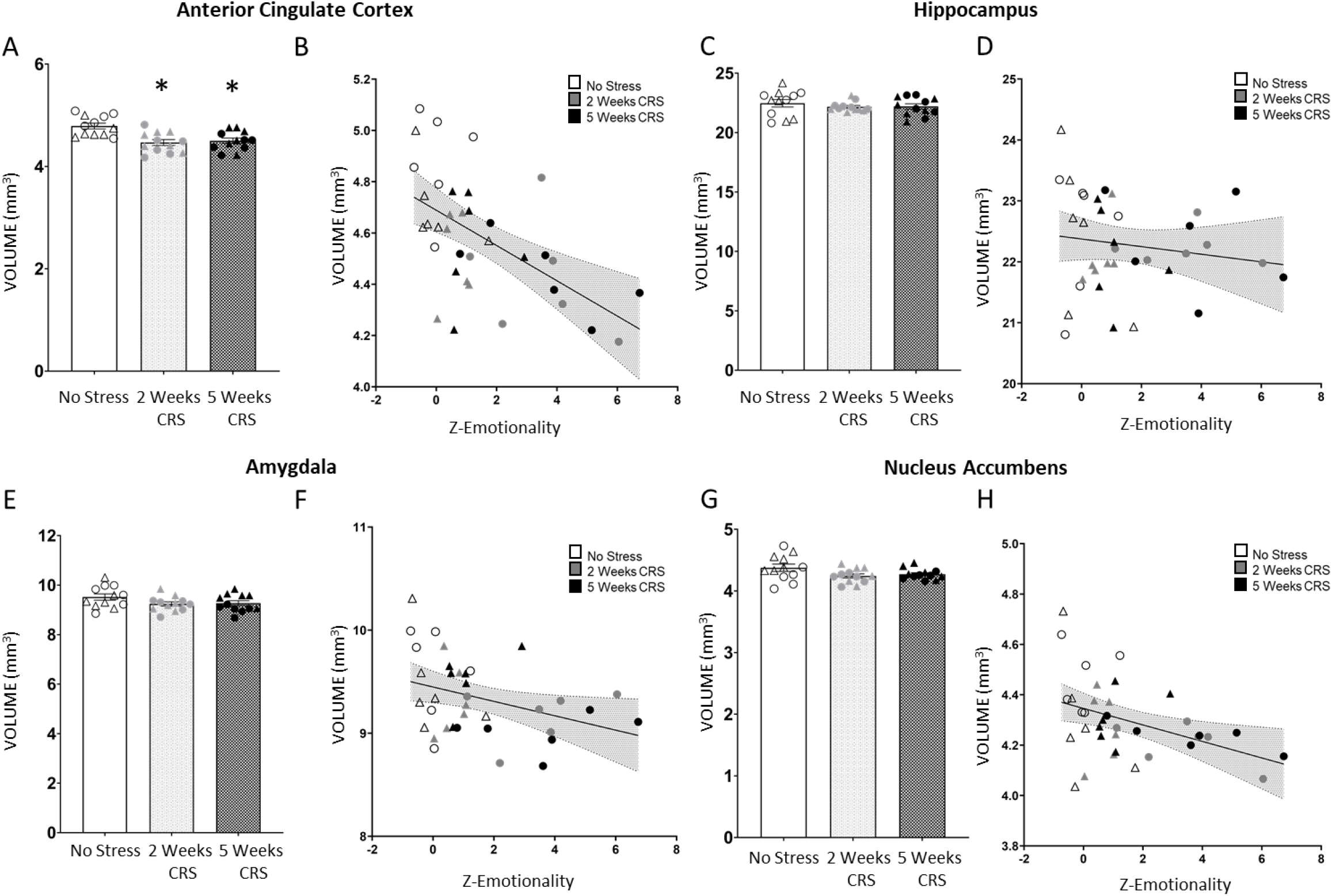
Chronic stress effects on 4 corticolimbic brain region volumes and correlation between regional volume and behavioral emotionality. Chronic restraint stress (CRS) induced a decreased in absolute volume of the (A) anterior cingulate cortex (*p<0.05 as compared to controls). No significant changes in absolute volume were found in the (C) hippocampus, (E) amygdala and (G) nucleus accumbens. Scatter plots display correlations (95% confidence interval) between z-emotionality score and volume changes in the (B) anterior cingulate cortex (r=-0.56; p=0.0003), (F) amygdala (r= −0.35; p=0.03), and (H) nucleus accumbens (r=-0.41; p= 0.01). No significant correlation observed in the (D) hippocampus. Individual males (Δ) and females (O) are represented in each figure.

### 3.4 CRS alters volumes of 2 of the 26 ROIs associated with MDD and 9 negatively correlated with z-emotionality

A follow-up analysis was performed across 26 ROIs involved in MDD **(Supplementary Table 1)** selected as in Nikolova et al. (2018). ANCOVA analysis using TBV as covariate and sex as cofactor revealed that the cingulate cortex area 24B and the striatum showed significant reductions in volume in the 2 or 5 weeks CRS groups as compared to controls after FDR correction for multiple comparisons (q<0.05). Pearson correlations were performed on the 26 ROI volumes corrected for TBV revealed a significant negative correlation between z-emotionality and volume of the cingulate cortex area 24A, 24B, hypothalamus, insular region, medial parietal association cortex, midbrain, NAc, striatum and thalamus (q<0.05, **Supplementary Table 2**). Logistic regression analysis was then conducted to determine association between 26 ROIs and Y-maze performance and revealed significant links with cingulate cortex area 29c, temporal association area and thalamus (p<0.05, **Supplementary Table 3**); however, these changes did not survive FDR correction. A follow-up whole-brain regional volume analysis of 182 brain regions was performed in all animals where only the primary somatosensory cortex: trunk region volume was significantly reduced in the 2 weeks CRS group compared to controls (q<0.05). The same analysis performed only in females showed that none of the 182 ROI display significant volumetric changes following CRS (all q>0.05) (**Supplementary Table 4 and 5**).

### 3.5 Progressive decrease in structural covariance network (SCN) degree in ACC and increase in SCN degree in the Amygdala in CRS exposed animals

SCN analysis can identify synchronous structural changes based on mutual trophic effects or shared plasticity between regions and networks (Mechelli, 2005). Additional recent work suggests structural covariance patterns further correspond to inter-regional structural connectivity and transcriptomic similarity (Yee et al., 2018). For the four selected corticolimbic brain regions, degree centrality was assessed to identify their position and “hubness”, respective to brain wide SCN changes. We found no significant group differences in regional degree or degree rank when comparing (1) the control group vs. 2 weeks CRS group or (2) the 2 weeks CRS group vs. 5 weeks CRS group. When comparing the control group and the 5 weeks CRS group, we found no significant group differences in degree or degree rank of the HPC or NAc (data not shown). However, comparison between the control and 5 weeks CRS groups for the ACC revealed that degree was significantly lower at multiple consecutive density thresholds in the 5 weeks CRS group compared to controls (density = 0.13 - 0.19, p ≤ 0.05, **Figure 4A**), with group differences largest at density = 0.16 (p = 0.012). Correspondingly, ACC degree rank was significantly lower in mice subjected to 5 weeks CRS compared to controls (density = 0.12 - 0.18, p ≤ 0.05). Using the first significant density threshold, the ACC SCN neighboring nodes (density = 0.12) are represented in **Figure 4B** for all three groups (see **Supplementary Table 6** for additional details). When comparing the control group and the 5 weeks CRS group, we found significant group differences in AMY degree whereby degree was significantly greater in the 5 weeks CRS group than in the control group at several consecutive density thresholds (density = 0.08 – 0.12, p ≤ 0.05, **Figure 4C**). Group differences were largest at density = 0.08 (p = 0.001). Amygdala degree rank was significantly lower in the 5 weeks CRS group than in the control group at several consecutive density thresholds (density = 0.05 - 0.10, p ≤ 0.05). Structural covariance networks for the AMY are represented in **Figure 4D,** where they are visualized at the first of the five aforementioned consecutive significant density thresholds (density= 0.08). Supplementary **Table 7** indicates neighboring nodes, the volume of which covaries with that of the AMY at the chosen density threshold. Connected and unconnected nodes are indicated with 1 and 0, respectively. For the AMY, we confirmed increased SCN degree and degree rank in animals subjected to chronic stress (Nikolova et al., 2018).

**Figure 4:**
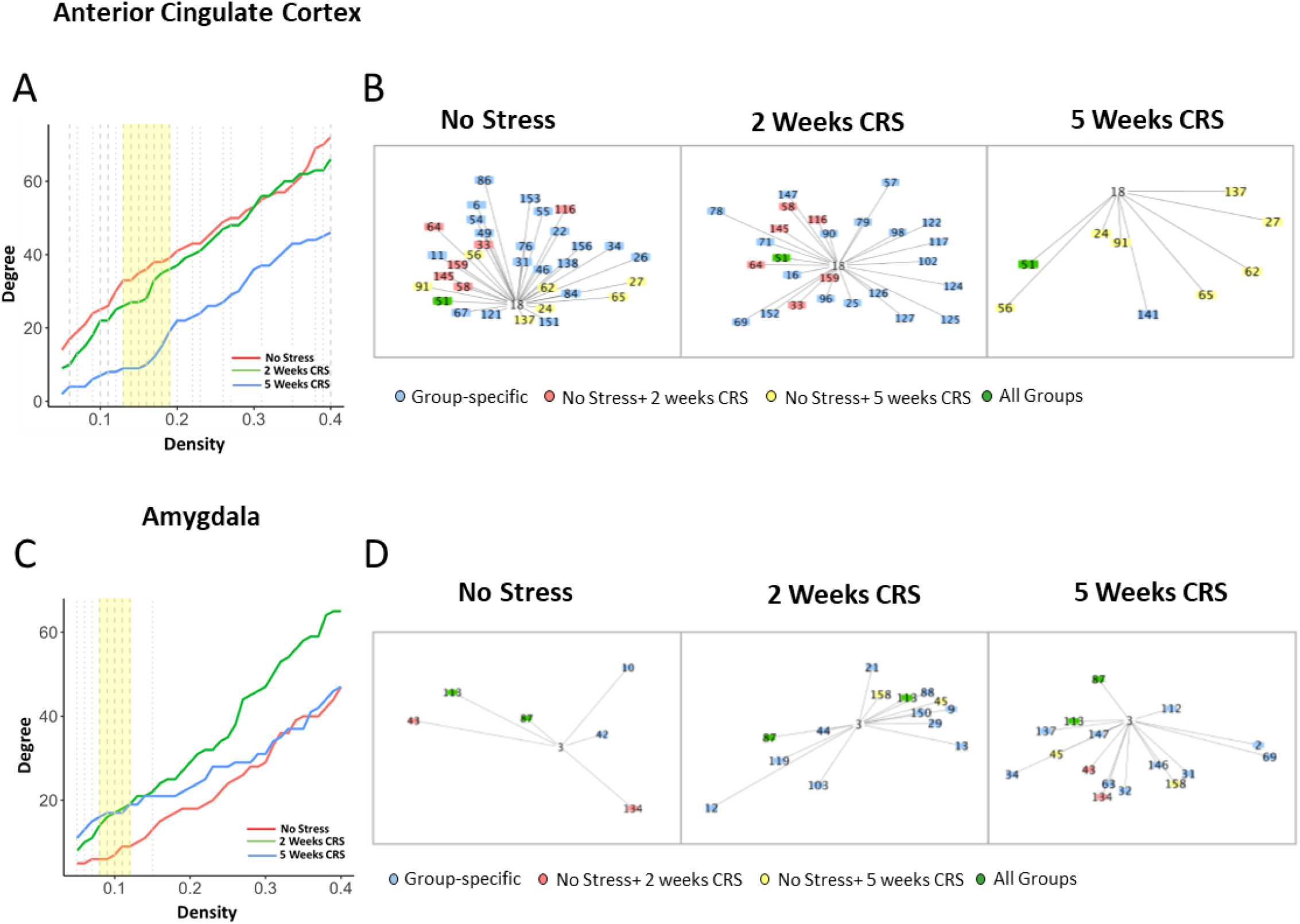
Chronic stress induced progressive decrease in strength and degree of the anterior cingulate cortex (ACC) and increase in strength and degree of the amygdala structural covariance networks. Decrease in ACC degree (A) were found in mice exposed to 2 and 5 weeks of chronic restraint stress (CRS), relative to mice exposed to no stress (controls). Permutation testing revealed significant differences in degree when comparing the 5-weeks CRS group to the control group at density = 0.13 – 0.19 (yellow) (p<.05 dashed, p<.10 (dotted)). Subnetwork visualization (B) at density = 0.13 was performed using the Cytoscape Edge-Weighted Spring Embedded Layout. Increase in amygdala degree (C) were found in mice exposed to 2 and 5 weeks chronic restraint stress (CRS), relative to mice exposed to no stress (controls). Permutation testing revealed significant differences in degree when comparing the 5week CRS group to the control group at density = 0.08 – 0.12 (yellow) (p<.05 dashed, p<.10 (dotted)). Subnetwork visualization (D) at density = 0.08 was used. Nodes labeled in red are common to the no stress and 2 weeks CRS groups; nodes labeled in yellow are common to the no stress and 5 weeks CRS groups; nodes labeled in green are common to all three stress groups; and nodes labeled in blue are unique to each stress group. Node numbers represent brain regions listed in **Supplementary Table 6 (ACC) and 7 (Amygdala)**. The length of each edge represents the strength of correlations between nodes.

### 3.6 PSD95 puncta density in the ACC correlates with ACC volume and z-emotionality

Given the significant effect of CRS exposure on the ACC volume, we investigated the changes in density of marker expression of pre- and post-synaptic compartments, PSD95 (**Figure 5A-C**) and VGLUT1 **(Supplementary Figure 6),** respectively. Quantitative analysis of PSD95 puncta density using analytic IMARIS quantitative spot counting software revealed a near-trend toward a main effect of stress (F_(2,28)_ =2.408; p=0.11), significant main effect of sex (F_(1,28)_ =6.462; p<0.05) and no stress x sex interaction (**Figure 5D**). Overall females displayed lower PSD95 puncta density but no significant differences in density of puncta between groups was found (data not shown). Interestingly, we found a significant positive correlation between PSD95 puncta density with volume changes of the ACC (*r*=0.35; p=0.044; **Figure 5E**). In addition, PSD95 puncta density was negatively correlated with z-emotionality score (*r*=-0.439; p=0.009; **Figure 5F**). Similar analysis conducted on VGLUT1 puncta density in ACC revealed no significant effect of stress, sex or stress x sex interaction, and no correlation with either z-emotionality or volume change of the ACC (**Supplementary Figure 6**). Using binary logistic regression, we found that PSD95 or VGLUT1 puncta densities did not predict Y-maze performances (χ^2^ = 1.62 and χ^2^ = −0.578, respectively). Given the significant effect of CRS exposure on SCN AMY, we also investigated the changes in PSD95 and VGLUT1 puncta density in the BLA and found no significant effects of stress or sex on PSD95 puncta density in this area. PSD95 puncta density in the BLA did not correlate with z-emotionality or AMY volume. However, CRS exposure induced an increase in VGLUT1 puncta density (p<0.05; **Supplementary Figure 7**) which correlated with z-emotionality (*r*=0.362, p<0.05) but not with AMY volume (**Supplementary Figure 7**).

**Figure 5:**
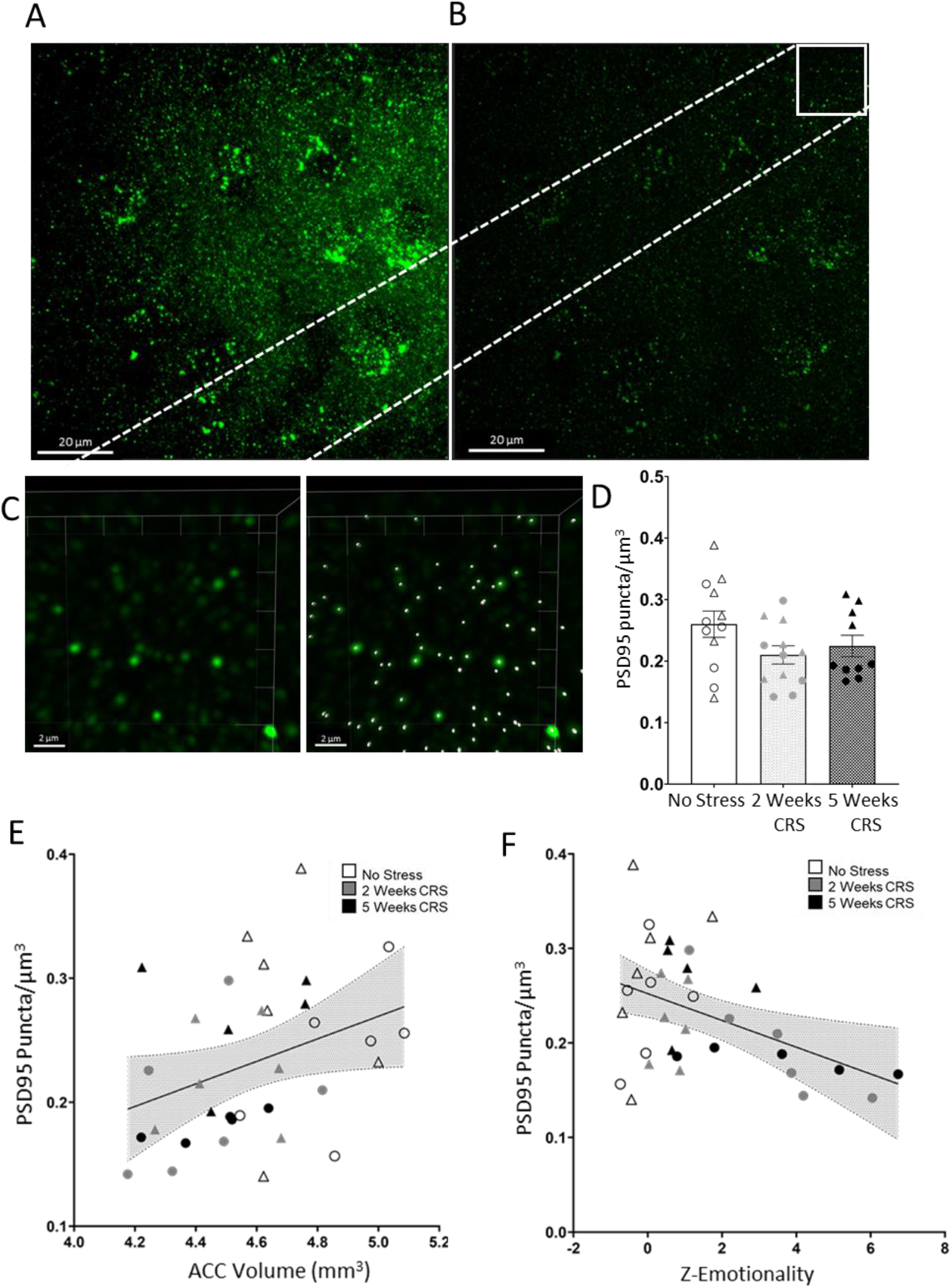
Changes in PSD95 puncta density are linked to volume of the anterior cingulate cortex (ACC) and behavioral emotionality. Max projection of three-dimension (3D) image z-stacks of top ~4μm of tissue imaged at 0.1μm/stack of (A) original confocal image of PSD95 immunolabelling and (B) blind deconvoluted image. (C) Image quantification performed using IMARIS software identifies quality of detected puncta based on intensity at center of each spot in 3D. CRS did not induce any significant changes in PSD95 puncta density of mouse anterior cingulate cortex (D). Scatter plots display a significant correlation (95% confidence interval) of changes in PSD95 puncta density with (E) ACC volume (r=0.35; p=0.044), and (F) behavioral emotionality (r=-0.439; p=0.009). Individual males (Δ) and females (O) are represented in each figure.

## 4 Discussion

The goal of this study was to use high resolution structural neuroimaging in rodents for studying brain-network adaptations underlying the behavioral deficits associated with chronic stress exposure. We showed that mice exposed to CRS displayed a progressive increase in anxiety-like behavior in the NSF and PhenoTyper tests, anhedonia-like behavior in the sucrose consumption test and impairment in working memory in the Y-maze test. These results confirmed that CRS results in increased behavioral emotionality and cognitive dysfunctions. Using ex-vivo MRI, we were able to identify an overall decrease in TBV (driven by reduced TBV in female mice) exposed to CRS. Correlation analyses revealed that reduced TBV was associated with increase in behavioral emotionality across sexes. Based on the literature in both animal and human studies of chronic stress and MDD, the ACC, AMY, NAc, and HPC were selected for our *a priori* analysis to identify changes in volume of corticolimbic structures. This analysis identified that CRS induced a significant decrease in volume of the ACC – a region of the mPFC commonly affected by chronic stress. Although no significant changes were observed in the other *a priori* corticolimbic brain regions, we were able to identify a link between the ACC, AMY and NAc volumes and behavioral emotionality deficits associated with CRS. We also confirmed that CRS-exposed mice show an increase in structural covariance degree of the AMY coupled with a decrease in ACC degree following 5 weeks of CRS exposure. The volumetric changes observed in the ACC were associated with PSD95 but not VGLUT1 puncta density. Additionally, reduced PSD95 puncta density in the ACC was negatively correlated with increased behavioral emotionality and positively correlated with reduced volumetric changes of this brain region. Finally, although CRS induced impairment in the Y-maze, performances in this test were not predicted by volumetric changes in specific brain region or synaptic puncta density in ACC. Altogether, chronic stress induced ACC volume reduction which was associated with reduced ACC structural covariance degree, PSD95 puncta density and increased behavioral emotionality, suggesting that altered synaptic strength is an underlying substrate of the volumetric and behavioral effects of CRS. Our results highlight a vulnerability of the ACC to chronic stress exposure and further implicate the ACC morphological alterations in stress-related illness including mood and anxiety disorders.

In accordance with the literature, our study confirmed that CRS induced behavioral emotionality in tests measuring anxiety- (Guilloux et al., 2011; Prevot et al., 2019b) and anhedonia-like behaviors (Banasr et al., 2010) as well as deficits in working memory (Prevot et al., 2019a). Prior work has shown that the effects of chronic stress are heterogeneous across sexes (Guilloux et al., 2011; Seney and Sibille, 2014). Here, we confirmed that female mice showed a progressive and higher magnitude of vulnerability to CRS exposure on anxiety- and anhedonia-like readouts compared to males (Guilloux et al., 2011). This is in accordance with human studies reporting greater prevalence of stress-induced symptoms of anxiety and comorbidity with anxiety disorders in women (Curry et al., 2014; Maeng and Milad, 2015; Schuch et al., 2014). These data critically emphasize the importance of using both sexes in experimental designs. Interestingly, in the Y-maze, a working memory task, deficits were found in male CRS mice. These results are in consensus with the literature describing deficits in the Y-maze appearing 7-10 days following CRS exposure (Prevot et al., 2019a). This result suggests that compared to male mice, females may exhibit lower susceptibility to cognitive impairments following chronic stress exposure. This hypothesis needs to be explicitly confirmed in future work but it is consistent with previous reports using other chronic stress rodent models (Conrad et al., 2003; Darcet et al., 2016). The progressive effects of chronic stress exposure were also confirmed on physical readouts such as degradation of coat state quality and lack of weight gain (Nikolova et al., 2018; Prevot et al., 2019b). Overall, regardless of sex differences our results identify behavioral emotionality deficits and cognitive impairments due to chronic stress exposure.

We also identified sex-specific patterns of volumetric changes, which complement prior rodent *in vivo* and *ex-vivo* MRI studies performed in male-only cohorts (Anacker et al., 2016; Lee et al., 2009; Magalhães et al., 2018; Nikolova et al., 2018). Specifically, we found that CRS induced an overall decrease in TBV driven by the female groups. Although, one prior study has reported TBV reductions in male rats following exposure to chronic unpredictable stress (Magalhães et al., 2018), other studies in male rodents have not reported such an effect (Lee et al., 2009; Nikolova et al., 2018). Imaging studies have associated between-group differences in TBV with variations in weight (O’Brien et al., 2011). In our study we found that although weight was significantly reduced by CRS, the association between weight and TBV did not account for much shared variance (data not shown); thus, weight was not regressed out in our analysis. The lack of consistency across studies regarding chronic stress-induced TBV changes in males may be due to the use of various chronic stress models, length of stress exposure or species (mouse or rat). Nevertheless, the female-specific reduction in TBV may also reflect greater susceptibility to chronic stress in female mice that may result in an overall brain atrophy. Studies have reported that intra-cranial volume was a more accurate readout than TBV to correct for, when studying conditions associated with neuronal atrophy (O’Brien et al., 2011). Interestingly, TBV reductions correlated with behavioral emotionality in both males and females. This suggests that a smaller TBV resulting from chronic stress might be an indicator of behavioral emotionality deficits. In human literature, total gray matter and TBV changes in MDD patients have been found but considered insignificant when compared to the general population (Zhuo et al., 2019). This overall brain reduction in rodents subjected to chronic stress may be the result of neuronal atrophy affecting numerous brain regions and being greater in females.

*A priori* analysis focusing on 4 corticolimbic brain regions identified the ACC as a brain region whose volume significantly decreased with CRS. We found a significant ~3% decrease in ACC volume in mice exposed to either 2 or 5 weeks CRS. This confirms previous MRI findings describing decreased ACC volume in C57Bl/6 mice following CRS (Kassem et al., 2013). In this latter study, The authors report a 16% reduction of ACC volume following 21 days of CRS (1 session of 6h restraint per day) (Kassem et al., 2013). The difference in magnitude of change may be due to greater length of each CRS session (Kassem et al., 2013). From human MRI literature, ACC volume reductions have been identified in major depressive (Bora et al., 2012; Van Tol et al., 2010), anxiety (Van Tol et al., 2010), and panic disorders (Asami et al., 2008). This reduced volume was identified in MDD patients who have experienced multiple episodes (Bora et al., 2012), lifetime occurrence of MDD (Ancelin et al., 2019), and are seen in treatment non-responders (Liu et al., 2017). Regarding the HPC, contrary to previous studies describing decreased volume following CRS (Kassem et al., 2013; Lee et al., 2009), we found no HPC volume change. This may be due to differences in length or duration of the CRS procedures or the fact that we did not subdivide the dorsal hippocampus from the ventral hippocampus, the latter known to be more involved in emotion regulation and more susceptible to stress (Kheirbek and Hen, 2011). We also found no significant changes in absolute volumes of the AMY and NAc volume in CRS mice, confirming most rodent chronic stress MRI studies (Anacker et al., 2016; Kassem et al., 2013; Lee et al., 2009; Magalhães et al., 2018). However, we previously described a ~3% increase in AMY relative volume following unpredictable chronic mild stress in BalbC male mice (Nikolova et al., 2018). Such inconsistencies in rodent chronic stress models mirror heterogeneous findings reporting variable results in human MDD patients with some studies describing larger, smaller or no changes in amygdala volume. In fact, smaller AMY volume has been attributed to greater number of episodes (Frodl et al., 2002; Frodl et al., 2003; Hamilton et al., 2008). Overall, in our study, we see volumetric changes in corticolimbic regions that may reflect a later state of the chronic stress response as they are somewhat similar to the volumetric changes reported with multiple episodes of MDD. Despite not finding significant changes in absolute volumes of corticolimbic brain regions (except for the ACC), we did identify strong correlations between volume of the AMY and NAc, and behavioral emotionality. Extending this first *a priori* analysis to MDD-associated brain regions allowed us to uncover additional brain regions of relevance to the expression of behavior emotionality deficits including multiple medial prefrontal subfields and key subcortical brain regions involved in sensory and emotion processing (thalamus, hypothalamus and midbrain). These correlations further emphasize the importance these regions have on the expression of depressive-like behavior and their role in the modulation of chronic stress response.

The seed-based analysis of structural covariance performed for the 4 corticolimbic brain regions identified that the ACC and the amygdala SCN connectivity patterns are affected by chronic stress. Specifically, we have identified a progressive CRS-induced decrease in the number of brain regions that structurally covaries with the ACC (i.e. the SCN degree of the ACC). Interestingly, this CRS-induced decrease in ACC degree is associated with a loss of connectivity with critical brain regions such as the NAc, medial amygdala, thalamus, bed nucleus of the stria terminalis (BNST) and multiple cortical subregions following 5 weeks CRS exposure. Human imaging studies showed that the medial prefrontal cortical network, including the medial orbital-frontal cortex, thalamus and limbic basal ganglia, is important in the processing of emotionally salient information (Price and Drevets, 2010). A reduced SCN degree could reflect a disconnect of the ACC of its network which may be responsible for, or at the very least linked with, elevated behavioral emotionality deficits found following CRS. Despite no significant AMY volumetric changes, we confirmed an increase in AMY SCN degree and increased synaptic puncta density with greater chronic stress exposure (Nikolova et al., 2018). Similar increase in AMY SCN degree was also found in a human cohort with history of childhood trauma (Nikolova et al., 2018). Our results suggest that the AMY SCN changes are conserved across stress models, sexes and strains as well as across species. The ACC and AMY SCN reorganization patterns suggest a decrease in ACC SCN connectivity coupled with an AMY SCN strengthening which maps onto synaptic puncta density in the ACC and increased density in the BLA respectively. Interestingly synaptic changes in these brain regions parallels SCN reorganization changes that may collectively be involved in the impaired emotional regulation associated with chronic exposure to stress.

One of the most common factors associated with brain volume changes is synaptic and dendritic reorganization. Here, we found a marginal decrease in PSD95 and no change in VGLUT1 puncta density in the ACC in CRS exposed mice. Decreases in PSD95 protein in the ACC have been identified in rodents subjected to various chronic stress models (Kang et al., 2012; Li et al., 2011; Qiao et al., 2016; Son et al., 2012). In the present study, PSD95 puncta density in the ACC correlated with volumetric change in the ACC, i.e., decreased volume is associated with decreases in the density of PSD95 puncta. Regarding VGLUT1 we found no significant changes in puncta density in the ACC. Human and rodent studies have identified decreases in mRNA expression of VGLUT1 (Uezato et al., 2009; Zink et al., 2010), analysis of protein expression following chronic stress exposure in the cortical brain regions have identified mixed results (Farley et al., 2012; Garcia-Garcia et al., 2009; Yu et al., 2018). Changes in VGLUT1 puncta did not correlate with volume changes in the ACC in this study. This lack of difference in ACC VGLUT1 expression in chronic stress mice compared to controls may be due to the fact that VGLUT1 protein can originate from other brain regions that project to the PFC unaffected by CRS (Collins et al., 2018; McGarry and Carter, 2017). Unlike VGLUT1, expression of PSD95 protein is postsynaptic, involved in synaptic strength and expressed by neurons endogenous to ACC which could explain why PSD95 puncta density was significantly correlated with the ACC volume changes. Since PSD95 puncta density changes may index altered synaptic density and strength (Meyer et al., 2014), our results suggest that these synaptic dysfunctions are underlying substrates for the reduced ACC volume and decreased ACC SCN associated with CRS, together responsible for behavioral emotionality deficits.

This study has several limitations. First, while we were able to detect some sex-specificity in the behavioral and structural changes induced by CRS, our study was not originally designed to examine sex differences and was underpowered to specifically investigate volumetric changes within each sex. A challenge in this study was that most of the CRS effects did not survive correction for TBV because of the unexpectedly reduced TBV in CRS. The large shared variance between TBV and a majority of brain regions and the CRS-induced TBV impacted our ability to detect region-specific changes due to CRS when TBV was added as a covariate. Perhaps controlling for intracranial volume when neuronal atrophy is expected could address this limitation, as with some imaging studies focusing on neurodegenerative disorders (O’Brien et al., 2011). Second, high resolution *ex-vivo* MRI can delineate relatively small brain structures (182 in this study) which can be beneficial to understand the involvement of certain sub-structures but adds difficulty to the analysis where very few changes are able to survive FDR correction. Although it might be beneficial in some rodent MRI studies to have a very precise atlas that can detect minute volumetric changes to small brain regions, such parcellation may impede the comparison of our findings to human MDD studies. Indeed, in many human MRI studies regional brain volume structures are not subdivided as extensively as in rodents and having a harmonized atlas between human and rodent MRI studies could bridge this gap. Nonetheless, we present results from whole-brain volumetric analyses including all regions in this refined parcellation scheme in order to foster discovery science and hypothesis generation for future studies. Third, in this study we were unable to establish a link between cognitive impairment and morphological (structural or synaptic) changes. This is perhaps due to the use of a single behavioral test assessing one cognitive dimension (working memory); further studies employing a more comprehensive behavioral assessment of cognition are needed to address this specific question. Despite these limitations, we were able to demonstrate that ACC volumetric changes induced by chronic stress correlated with behavioral changes and PSD95 puncta density, linking these macroscale and microscale alterations to behavioral deficits. Altogether, our results highlight a vulnerability of the ACC to chronic stress at multiple levels (volume, connectivity and spine) and its role in the expression of emotionality in behaviors relevant to stress-related illness including mood and anxiety disorders.

## 5 Acknowledgements

This work was supported by the Canadian Institute of Heath research (CIHR Project Grant PJT#165852, PI: MB), the CAMH Discovery Fund (MB) and the Campbell Family Mental Health Research Institute of CAMH (MB, ES). YSN is supported by a Koerner New Scientist Award administered by the CAMH Foundation and an NSERC Discovery Grant (RGPIN-2020-07131). AM is supported by a CAMH Discovery Fund Postdoctoral Fellowship. JPL and JE are supported by the Ontario Brain Institute and CIHR.

## 6 Conflicts of Interests

The authors declare that they have no conflict of interest related with this work.

## Notes

### Competing Interest Statement

The authors have declared no competing interest.

