## Supplement Material for "Reduced anterior cingulate cortex volume induced by chronic stress correlates with increased behavioral emotionality and decreased synaptic puncta density"

### Table content

**Supplementary Results (p2-4)**

**Supplementary Figure 1 (p5-6)**

**Supplementary Figure 2 (p7)**

**Supplementary Figure 3 (p8)**

**Supplementary Figure 4 (p9)**

**Supplementary Figure 5 (p10-11)**

**Supplementary Figure 6 (p12)**

**Supplementary Table 1 (p13)**

**Supplementary Table 2 (p14)**

**Supplementary Table 3 (p15)**

**Supplementary Table 4 (p16-20)**

**Supplementary Table 5 (p21-26)**

**Supplementary Table 6 (p27-29)**

**Supplementary Table 7 (p30-31)**

### Supplemental Methods

All procedures were performed in accordance with institutional Animal Care Committee (ACC) and Canadian Council on Animal Care (CCAC) procedural and ethical guidelines. CAMH ACC guideline for humane end-points include: >20% weight loss, inability to attend to bodily needs, extremely hunched posture, or complete loss of fur, signs of distress such as vocalization, shivering, seizure or lethargy. However, none of the CRS animals of this study showed these signs. On week 2 of behavioral testing in the PhenoTyper test, the data for the one control female mouse was missing due a technical problem with the infrared-light tracking. We decided to input the missing data for that week for this mouse to allow us to not reduce the already relatively small sample size. For the other weeks, all data and all other behaviors were collected in their entirety for that mouse and all the other subjects. Only the data collected at week 5 were used for the behavior/ volume, behavior/synaptic puncta correlations. This did not affect any of the key findings of the manuscript.

### Supplemental Results

*Chronic stress exposure induces longitudinal and progressive changes in anxiety-like behaviour measured using the PhenoTyper test*

The PhenoTyper apparatus allows for monitoring time spent in the shelter zone (before, during and after a light challenge applied over the food zone) weekly over the course of the 5 weeks experimental procedure. Data were collected consistently over the course of the experiment except for one female mouse at the week 5 time point. Missing data were due to tracking acquisition issues and data for this one female mouse was imputed based on the mean of the group. Without splitting for sex, repeated measures ANCOVA reveals a significant main effect of stress on week 1 ($F_{(2, 360)} = 8.823$; $p < 0.01$), and every following week (weeks 2 ($F_{(2, 360)} = 11.144$; $p < 0.001$), 3 ($F_{(2, 360)} = 19.654$; $p < 0.001$), 4 ($F_{(2, 360)} = 27.159$; $p < 0.001$) and 5 ($F_{(2, 360)} = 14.424$; $p < 0.0001$). Specifically, mice exposed to 5 weeks of CRS (CRS5w) spend a progressive increased amount of time in the shelter between 12:00-05:00 for the first three weeks of CRS exposure as compared to controls and 2 weeks CRS group (CRS2w) before the start of the CRS. On Weeks 4 and 5, CRS5w and CRS2w animals showed increased time spent in the shelter as compared to controls specifically between 12:00-05:00 (**Supplementary Figure1B-G**). When sex is included as a cofactor in the analysis, a main effect of sex was found on weeks 3 ($F_{(1, 360)} = 6.610$; $p < 0.05$), 4 ($F_{(1, 360)} = 3.982$; $p = 0.0551$), and 5 ($F_{(1, 360)} = 6.847$; $p < 0.05$). A significant main effect of stress x time interaction on time spent in the shelter zone on week 1 ($F_{(24, 360)} = 1.639$; $p < 0.05$), week 2 ($F_{(24, 360)} = 3.491$; $p < 0.0001$), week 3 ($F_{(24, 360)} = 1.649$; $p < 0.05$), week 4 ($F_{(24, 360)} = 2.12$; $p < 0.01$), week 5 ($F_{(24, 360)} = 2.044$; $p < 0.01$) (**Supplementary Figure 1B-G**). Lastly a sex x time interaction was observed on week 5 ($F_{(12, 360)} = 1.786$; $p < 0.05$) and a stress x sex x time interaction only found on week 4 ($F_{(24, 360)} = 2.586$; $p < 0.0001$).

CRS exposed mice spend the majority of the time in the shelter zone after the light challenge between time points 12:00-05:00 (**Supplemental Figure 1**). This identified increase in time spent in the shelter zone was measured using the RA analysis. Repeated measures ANCOVA of residual avoidance in the shelter zone revealed significant overall main effect of stress ($F_{(2, 150)} = 13.849$; $p < 0.0001$; **Supplemental Figure 1H**) and stress x time interaction ($F_{(10, 150)} = 8.427$; $p < 0.0001$). Post-hoc analysis revealed a significant difference between controls and CRS5w animals on weeks 3, 4 and 5 ($p<0.01$) and between control and CRS2w animal groups on weeks 4 and 5 ($p<0.01$). A main effect of sex ($F_{(2, 150)} = 4.866$; $p < 0.05$; **Supplement Figure 4B**) stress x sex ($F_{(10, 150)} = 8.991$; $p < 0.0001$) and a stress x sex x time interaction ($F_{(10, 150)} = 3.766$; $p < 0.001$) (**Supplementary Figure 4B**) was found. Analysis within sex revealed a significant increase in RA in females during weeks 2,3,4, and 5 for the CRS5w mouse group and in weeks 4 and 5 in CRS2w group as compared to female controls. In male CRS5w mice exhibited an increase in RA in weeks 2,3,4, and 5 and in CRS2w mice an increase RA in weeks 4 and 5 as compared to male controls.

Corresponding behavioral output was observed in the food zone during the PhenoTyper test which present opposing effects to the shelter zone. Analysis of food zone time revealed a main effect of stress in weeks 3 ($F_{(2, 360)} = 13.936$; $p < 0.0001$), 4 ($F_{(2, 360)} = 11.494$; $p < 0.001$) and 5 ($F_{(2, 360)} = 3.84$; $p < 0.05$). We also found a significant stress x time interaction on week 2 ($F_{(24, 360)} = 1.901$; $p < 0.01$) and a trend towards significance on week 4 ($F_{(24, 360)} = 1.489$; $p = 0.06$). When sex is used a cofactor, we found a significant sex x time interaction at week 3 ($F_{(12, 360)} = 1.737$; $p < 0.05$) and 5 ($F_{(12, 360)} = 2.399$; $p < 0.01$). Analysis of the residual avoidance data in the food zone revealed a significant effect of stress ($F_{(2, 150)} = 4.468$; $p < 0.05$) due to a significant difference between controls or CRS2w animals and CRS5w group on weeks 2, 3, 4 and 5 ($p<0.01$) and between controls and CRS2w mice on weeks 4 and 5 ($p<0.01$). We found no significant effect of sex, stress x time and sex x time interaction found in the residual avoidance analysis of the food zone.

*Chronic stress exposure induces longitudinal and progressive changes in anhedonia-like behavior in the sucrose consumption test, coat state quality score and weight gain.*

Repeated measures ANCOVA of weekly sucrose consumption revealed no significant main effect of stress, sex (trend p=0.07), stress x time, stress x sex x time interaction (**Supplementary Figure 2B/4E**). Repeated measures ANCOVA indicated a significant main effect of stress ($F_{(2, 150)}$ = 7.352; p < 0.01), A *post-hoc* analysis revealed a significant difference in weight gain between CRS5w mice compared to controls in weeks 3 and 5 (p<0.01; **Supplementary Figure 2C).** A main effect of sex ($F_{(1, 150)}$ = 186.449; p < 0.0001), stress x time interaction ($F_{(10, 150)}$ = 10.23; p < 0.0001) in the assessment of weight gain. Analysis in female mice revealed a significant decrease in weight on weeks 2,3,4 for the CRS5w mice and in weeks 4 and 5 for the CRS2w mice. In male mice, a significant decrease was observed in week 1,3,4,5 for the CRS5w mice and in week 4 for CRS2w mice when compared to male controls (**Supplement Figure 4C**).

Repeated measures ANCOVA of coat state degradation revealed a significant main effect of stress ($F_{(2, 150)}$ = 31.192; p < 0.0001). Post-hoc analysis revealed a significant difference in coat state between CRS5w mice and with control or CRS2w mouse group in weeks 2, 3, 4, and 5 (**Supplementary Figure 2D**). A main effect of sex ($F_{(1, 150)}$ = 30.032; p < 0.0001), stress x time ($F_{(10,150)}$ = 3.869; p < 0.001) and sex x time ($F_{(10,150)}$ = 2.665; p < 0.05) interaction were found. Post-hoc analysis within sex revealed a significant increase in coat state score in CRS5w females on weeks 2 and 4. While CRS5w males displayed a significant increase in coat state score in weeks 2,3,4, and 5 (**Supplementary Figure 4D**).

*CRS mice reveal difference in VGLUT1 puncta in the ACC*

Analysis using IMARIS spot counting software VGLUT1 puncta density revealed no significant differences between control and either stress groups (Supplement Figure 7). Pearson correlational analysis revealed no significant correlation between VGLUT1 puncta density and volume changes in the ACC (**R=-0.034; p=0.849, Supplement Figure 6E**) or z-emotionality (**R=0.013; p=0.940, Supplement Figure 6F**).

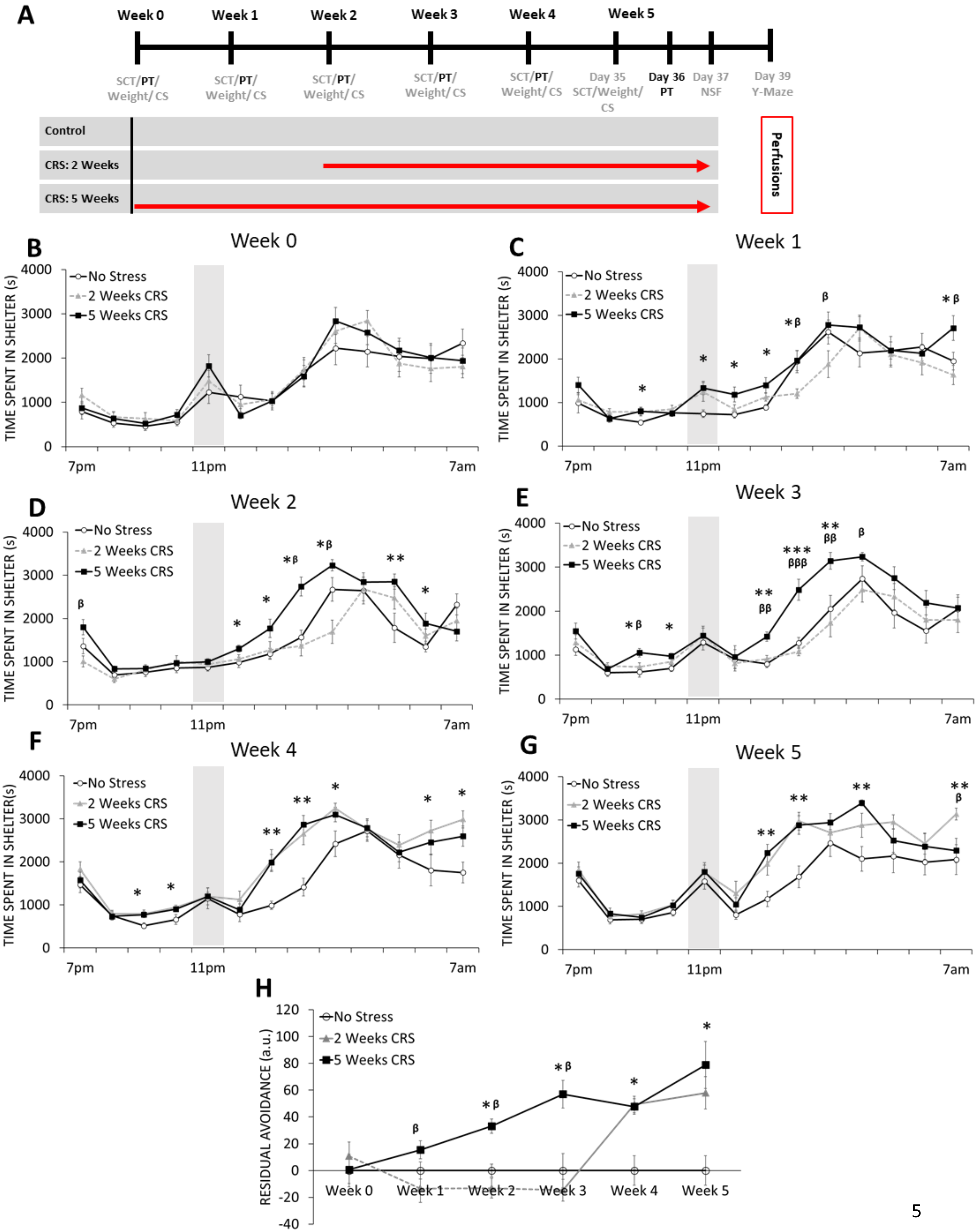
A
Week 0
Week 1
Week 2
Week 3
Week 4
Week 5
SCT/PT/Weight/CS
Day 35 SCT/Weight/CS
Day 36 PT
Day 37 NSF
Day 39 Y-Maze
Control
CRS: 2 Weeks
CRS: 5 Weeks
Perfusions
B
C
D
E
F
G
H
No Stress
2 Weeks CRS
5 Weeks CRS
TIME SPENT IN SHELTER (s)
7pm
11pm
7am
RESIDUAL AVOIDANCE (a.u.)

**Supplementary Figure 1: Longitudinal assessment in the PhenoTyper test revealed a progressive increase in anxiety-like behavior measured as residual avoidance of the food zone in favor of the shelter in mice exposed to 2 and 5 weeks of chronic stress.** A) Schematic representation of the experimental design. Control animals and mice subjected to 2 or 5 weeks of chronic restraint stress (CRS) (n=12/group; 50% males) were tested in series of behavioral tests throughout the experiment (gray). Mice were tested weekly in the PhenoTyper test (black) between 7PM to 7AM for a 5 week testing period. A white spotlight challenge occurs between 11-12PM (gray bars). (B) The time spent in the shelter zone at baseline (week 0) and (C-G) every following week was assessed. (H) Residual avoidance was calculated for each week and graphed to illustrate progression of avoidance behavior of the food zone in favor of the shelter over the course of the experiment. For B-E and H, we used doted lines for animal group subjected to CRS 2 weeks illustrating performance before the beginning of the CRS exposure and plain lines for performances recorded during CRS exposure (F-H). (*$p < 0.05$, **$p < 0.01$, ***$p<0.001$ as compared to controls and $^{\beta}p<0.05$, $^{\beta}p<0.01$ s compared with 2 weeks CRS)

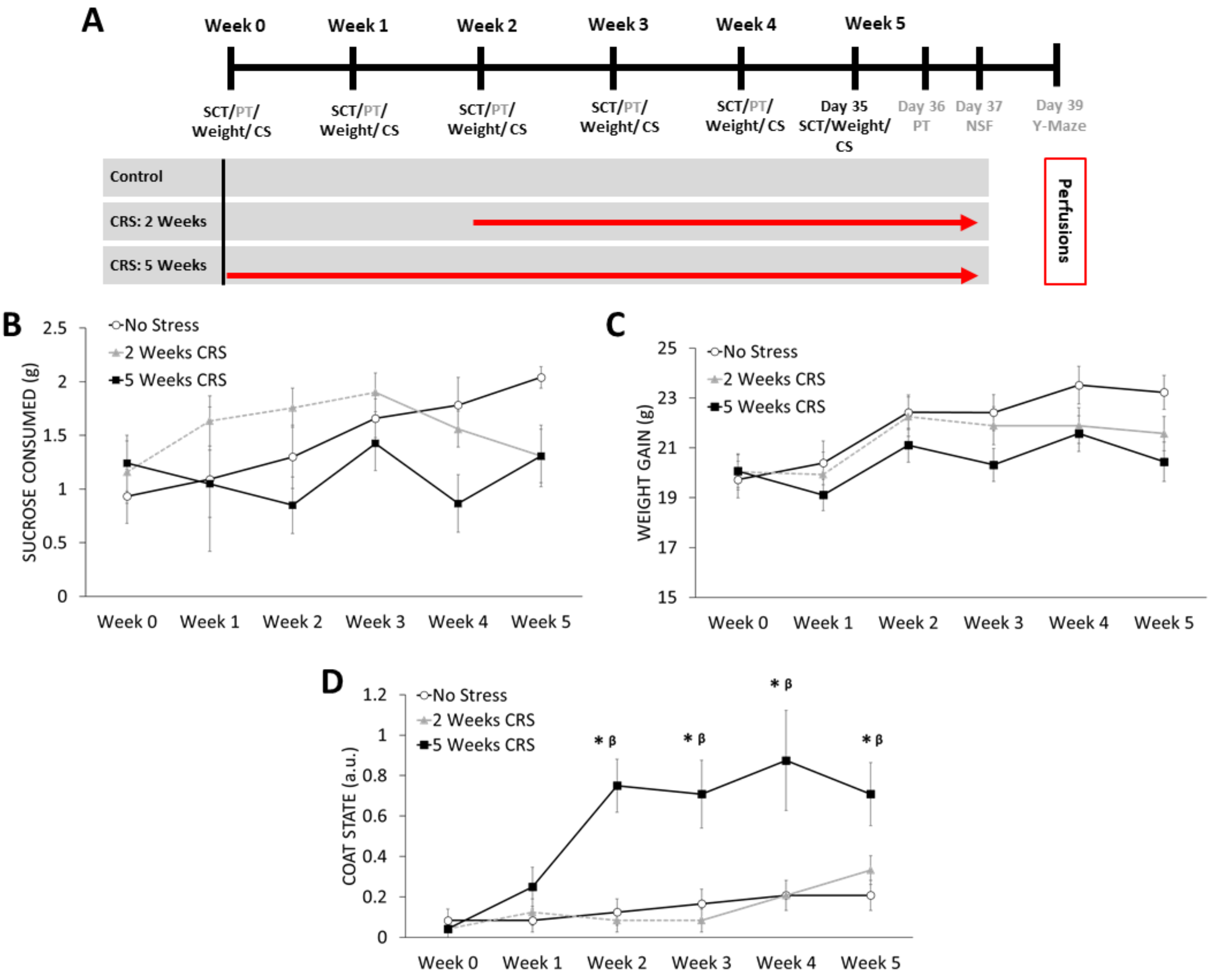


**Supplementary Figure 2: Longitudinal effects of chronic stress on weekly sucrose consumption, weight, and coat state assessments.** A) Schematic representation of the experimental design. Control animals and mice subjected to 2 or 5 weeks of chronic restraint stress (CRS) (n=12/group; 50% males) were tested in series of behavioral tests throughout the experiment (gray). Mice were tested weekly in the sucrose consumption test, weight and coat state assessment (black) during 5 weeks testing period. (B) sucrose consumption and (C) weight and (D) coat state degradation were measured in animals subjected or not to chronic restraint stress (CRS) (n=12/group). *P<0.05 compared with no stress; $^{\beta}$p<0.05 compared with 2 weeks CRS group. For the animal group subjected to CRS 2 weeks, doted lines were used for the illustrating the performances before the beginning of the CRS exposure and plain lines for the performances recorded during CRS exposure.

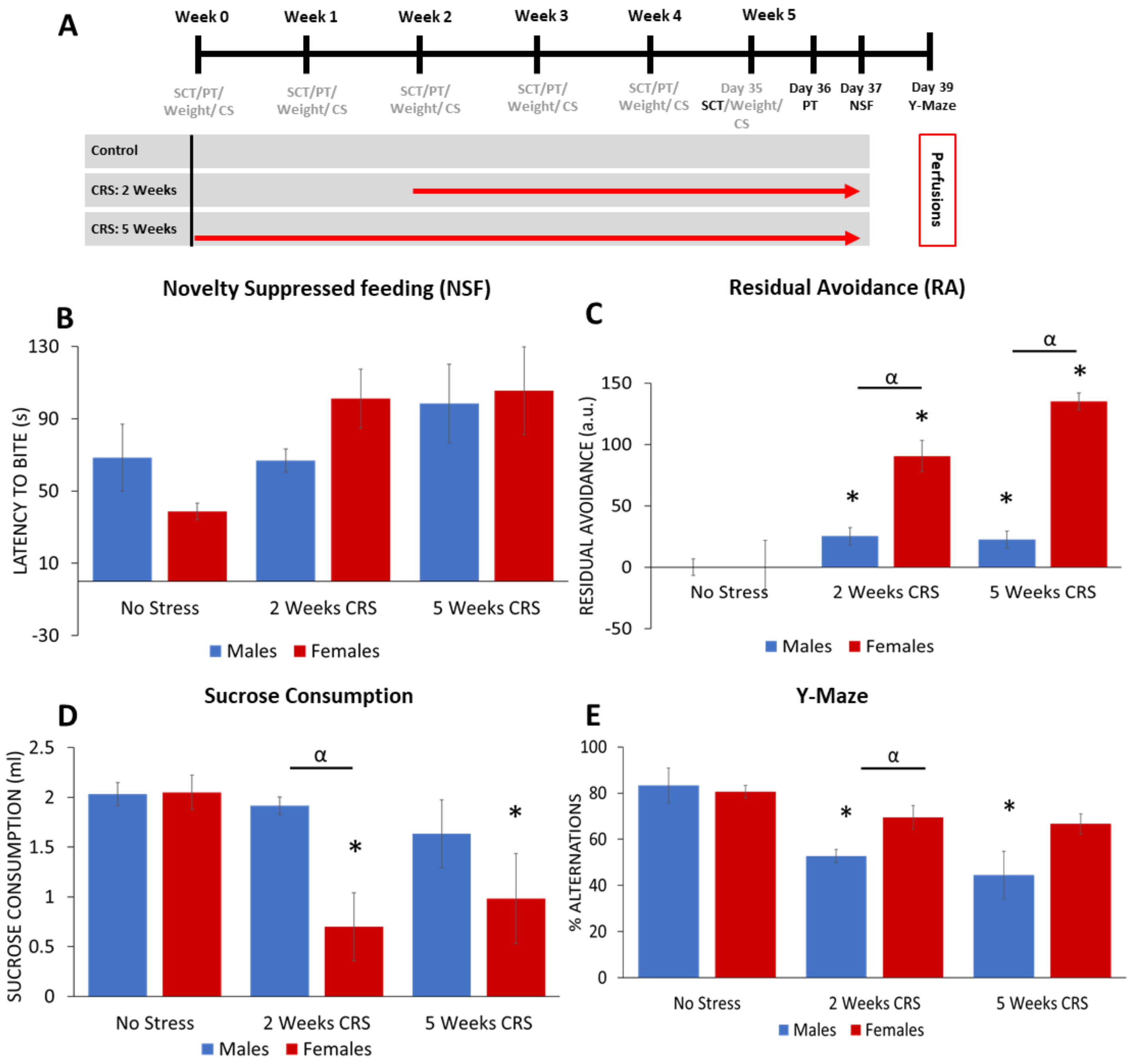


**Supplementary Figure 3: Differential effects of chronic restraint stress-induced deficits in behavioral emotionality and working memory across sexes.** (A) Schematic representation of the experimental design. Male and female C57Bl/6 mice (n=6/group) were subjected or not to chronic restraint stress (CRS) for 2 or 5 weeks across a series of behavioral tests throughout the experiment (gray). This figure summarized the data collected in the behavioral tests conducted during the 5$^{th}$ week of the experiment (black). Mice were tested in the (B) novelty suppressed feeding, (C) PhenoTyper, (D) sucrose consumption and (E) Y-maze tests. *$p<0.05$ as compared to respective controls; $^{\alpha}p<0.05$ as compared to males

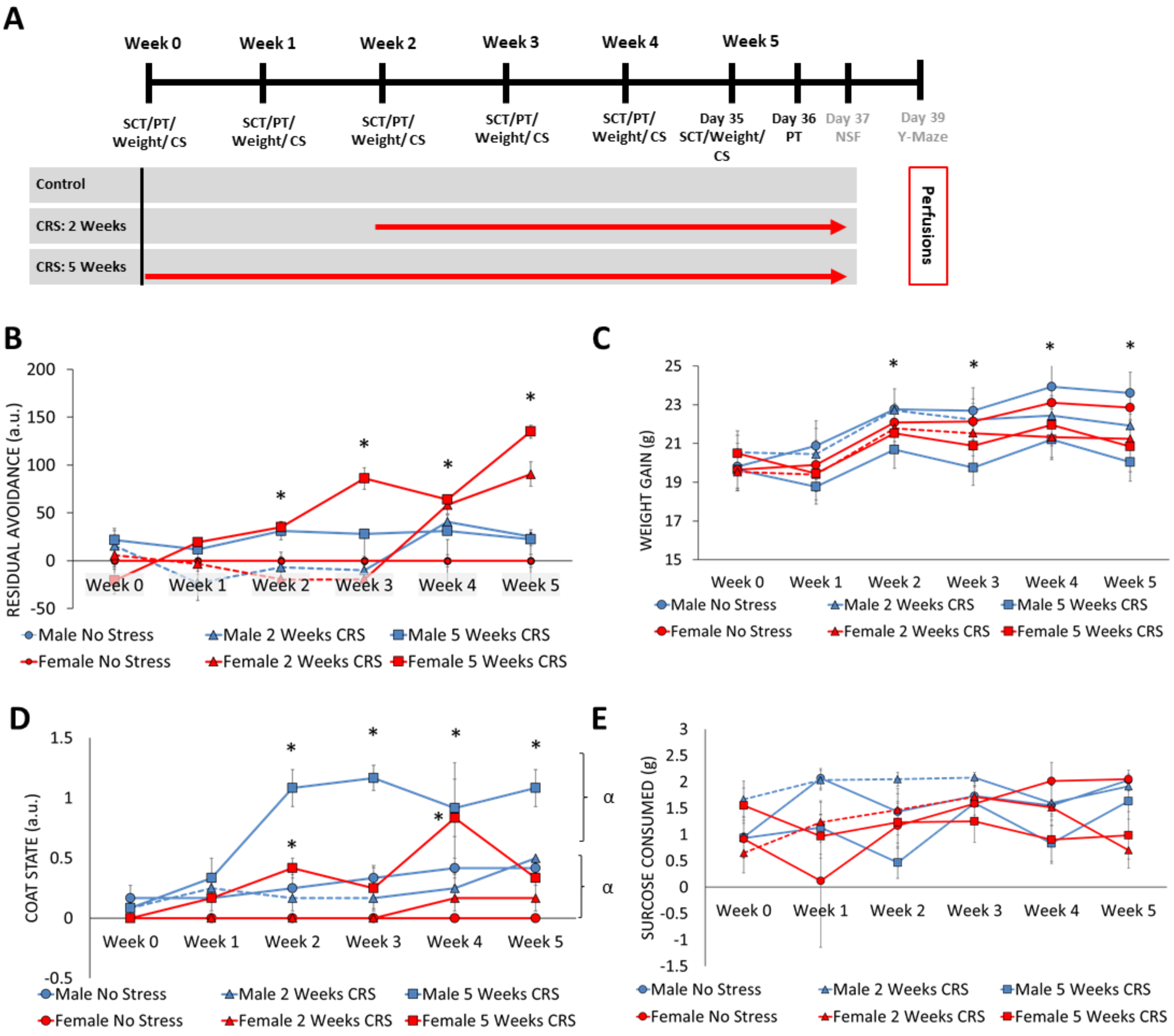


**Supplementary Figure 4: Differential effects of chronic restraint stress-induced deficits in weekly PhenoTyper, weight, and coat state assessments across sexes**. A) Schematic representation of the experimental design. Male and female C57Bl/6 mice (n=6/group) were subjected or not to chronic restraint stress (CRS) for 2 or 5 weeks across a series of behavioral tests throughout the experiment (gray). Mice were tested weekly in the PhenoTyper, sucrose consumption tests, weight and coat state assessment (black). Mice were assessed every week in the (B) PhenoTyper test, (C) weight, (D) coat state and (E) sucrose consumption. *$p<0.05$ as compared to respective controls; $^{\alpha}p<0.05$ as compared to males

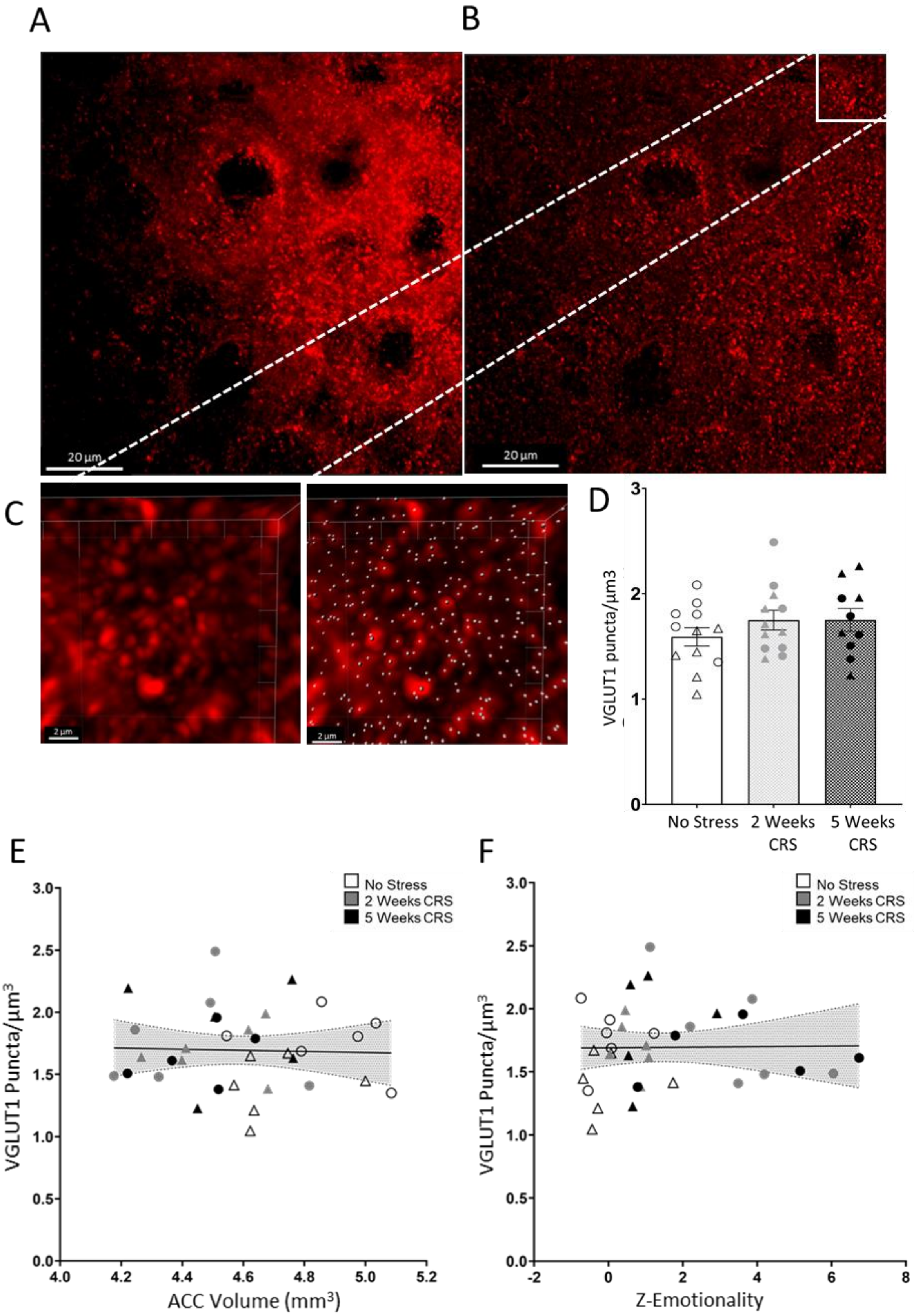
A
B
20 µm
20 µm
C
2 µm
2 µm
D
VGLUT1 puncta/µm3
No Stress
2 Weeks CRS
5 Weeks CRS
E
VGLUT1 Puncta/µm3
ACC Volume (mm3)
No Stress
2 Weeks CRS
5 Weeks CRS
F
VGLUT1 Puncta/µm3
Z-Emotionality
No Stress
2 Weeks CRS
5 Weeks CRS

**Supplementary Figure 5: VGLUT1 puncta density was not linked to volume of the anterior cingulate cortex (ACC) and behavioral emotionality.** Max projection of three-dimension (3D) image z-stacks of top ~4µm of tissue imaged at 0.1µm/stack of (A) original confocal image and (B) blind deconvoluted image. (C) Image quantification performed using IMARIS software identifies quality of detected spots based on intensity at center of each spot in 3D. CRS induced no significant changes in VGLUT1 puncta density of mouse anterior cingulate cortex (D). Scatter plots display no significant correlations (95% confidence interval) of changes in VGLUT1 puncta density with (E) anterior cingulate cortex volume and (F) behavioral emotionality indexed by the z-emotionality score. Individual males (△) and females (O) are represented in each figure.

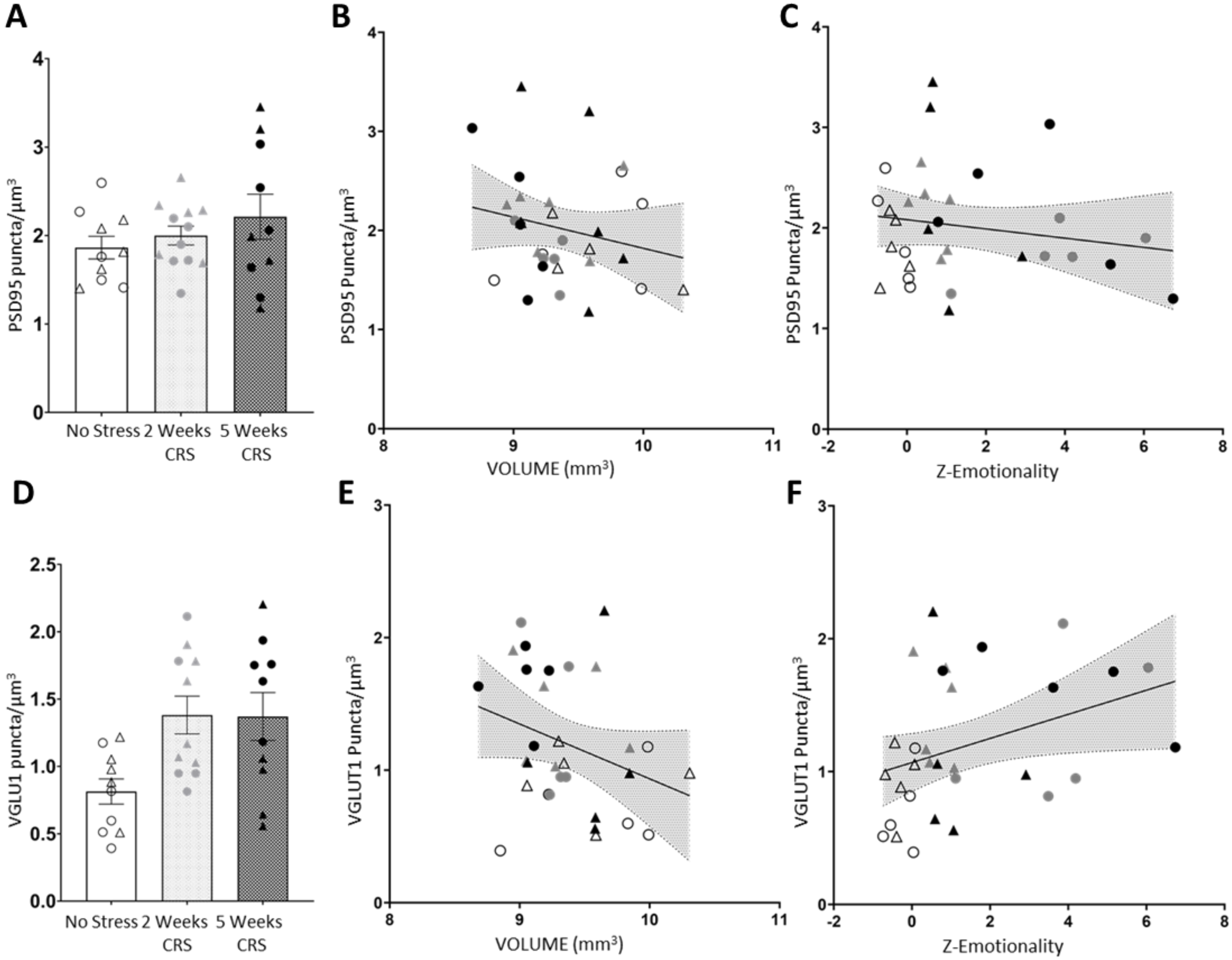


**Supplementary Figure 6: CRS induced altered synaptic puncta density in the basolateral amygdala (BLA).** (A) No significant changes in PSD95 puncta density in BLA was observed following CRS exposure. There was no correlation between PSD95 puncta density in BLA and (B) amygdala volume and (C) z-emotionality. (D) VGLUT1 puncta density was significantly increased by 2 and 5 weeks of exposure to CRS (* $p<0.05$). VGLUT1 density in BLA did not correlate with (E) amygdala volume but was significantly correlated with (F) z-emotionality. Individual males (△) and females (O) are represented in each figure.

**Supplementary Table 1**: Analysis of brain volumetric changes in 26 MDD-associated regions.

| ROI | p Value | q Value |
|---|---|---|
| **Cingulate cortex: area 24B** | **0.00** | **0.05** |
| **striatum** | **0.00** | **0.05** |
| Lateral parietal association cortex | 0.01 | 0.06 |
| Medial parietal association cortex | 0.02 | 0.14 |
| Frontal association cortex | 0.04 | 0.19 |
| Cingulate cortex: area 32 | 0.06 | 0.25 |
| thalamus | 0.07 | 0.25 |
| Cingulate cortex: area 29c | 0.23 | 0.65 |
| hypothalamus | 0.22 | 0.65 |
| Cingulate cortex: area 29a | 0.36 | 0.90 |
| Cingulate cortex: area 24A | 0.75 | 0.98 |
| Cingulate cortex: area 30 | 0.78 | 0.98 |
| Dorsolateral orbital cortex | 0.64 | 0.98 |
| globus pallidus | 0.64 | 0.98 |
| Insular region: not subdivided | 0.70 | 0.98 |
| Lateral orbital cortex | 0.76 | 0.98 |
| Medial orbital cortex | 0.72 | 0.98 |
| nucleus accumbens | 0.59 | 0.98 |
| Temporal association area | 0.53 | 0.98 |
| Ventral orbital cortex | 0.44 | 0.98 |
| amygdala | 0.88 | 0.98 |
| Cingulate cortex: area 25 | 0.92 | 0.98 |
| Cingulate cortex: area 29b | 0.94 | 0.98 |
| midbrain | 0.94 | 0.98 |
| Hippocampus | 0.99 | 0.99 |

**Supplementary Table 2**: Pearson correlation between 26 MDD-associated brain region volumes and z-emotionality.

| MDD Associated | Pearson R | p Value | q Value |
|---|---|---|---|
| **Hypothalamus** | -0.53 | 0.00 | 0.02 |
| **Striatum** | -0.49 | 0.00 | 0.02 |
| **Cingulate cortex: area 24B** | -0.48 | 0.00 | 0.02 |
| **Cingulate cortex: area 24A** | -0.47 | 0.00 | 0.02 |
| **Midbrain** | -0.47 | 0.00 | 0.02 |
| **Insular region: not subdivided** | -0.46 | 0.01 | 0.02 |
| **Nucleus accumbens** | -0.42 | 0.01 | 0.04 |
| **Thalamus** | -0.41 | 0.01 | 0.04 |
| **Medial parietal association cortex** | -0.40 | 0.01 | 0.04 |
| Globus pallidus | -0.38 | 0.02 | 0.06 |
| Amygdala | -0.36 | 0.03 | 0.07 |
| Temporal association area | -0.35 | 0.04 | 0.08 |
| Dentate gyrus of the Hippocampus | -0.32 | 0.06 | 0.11 |
| Lateral parietal association cortex | -0.28 | 0.10 | 0.18 |
| Dorsolateral orbital cortex | -0.28 | 0.10 | 0.18 |
| Lateral orbital cortex | -0.27 | 0.12 | 0.19 |
| Frontal association cortex | -0.26 | 0.12 | 0.19 |
| Ventral orbital cortex | -0.24 | 0.15 | 0.22 |
| Cingulate cortex: area 25 | -0.24 | 0.16 | 0.22 |
| Cingulate cortex: area 30 | -0.23 | 0.17 | 0.22 |
| Cingulate cortex: area 32 | -0.14 | 0.41 | 0.51 |
| Medial orbital cortex | -0.10 | 0.56 | 0.66 |
| Cingulate cortex: area 29c | -0.08 | 0.63 | 0.67 |
| Hippocampal Formation | -0.08 | 0.64 | 0.67 |
| Cingulate cortex: area 29b | -0.08 | 0.64 | 0.67 |
| Cingulate cortex: area 29a | -0.01 | 0.97 | 0.98 |

**Supplementary Table 3:** Binary logistic regression identifying associations between TBV-regressed ROI volumes and percent alternations in the Y-maze test.

| ROI | χ2 | p Value | q Value |
|---|---|---|---|
| **A priori** | | | |
| Amygdala | 1.50 | 0.13 | |
| Nucleus accumbens | -1.38 | 0.17 | |
| ACC | 1.71 | 0.09 | |
| Hippocampus formation | -0.62 | 0.53 | |
| **MDD Associated Regions** | | | |
| Temporal association area | -2.30 | 0.02 | 0.28 |
| Thalamus | 2.34 | 0.02 | 0.28 |
| Cingulate cortex: area 29a | 2.10 | 0.04 | 0.31 |
| Cingulate cortex: area 25 | -1.83 | 0.07 | 0.39 |
| Ventral orbital cortex | -1.78 | 0.08 | 0.39 |
| Medial parietal association cortex | -1.66 | 0.10 | 0.42 |
| Amygdala | 1.50 | 0.13 | 0.43 |
| Hippocampus | -1.56 | 0.12 | 0.43 |
| Nucleus accumbens | -1.38 | 0.17 | 0.44 |
| Striatum | 1.43 | 0.15 | 0.44 |
| Cingulate cortex: area 29c | -1.19 | 0.23 | 0.47 |
| Frontal association cortex | -1.11 | 0.27 | 0.47 |
| Globus pallidus | -1.26 | 0.21 | 0.47 |
| Hypothalamus | -1.13 | 0.26 | 0.47 |
| Insular region not subdivided | -1.10 | 0.27 | 0.47 |
| Lateral orbital cortex | -1.02 | 0.31 | 0.47 |
| Medial orbital cortex | -1.05 | 0.29 | 0.47 |
| Cingulate cortex: area 30 | -0.84 | 0.40 | 0.57 |
| Dorsolateral orbital cortex | -0.81 | 0.42 | 0.57 |
| Cingulate cortex: area 24A | -0.64 | 0.52 | 0.62 |
| Cingulate cortex: area 29b | -0.60 | 0.55 | 0.62 |
| Lateral parietal association cortex | -0.65 | 0.51 | 0.62 |
| Midbrain | 0.63 | 0.53 | 0.62 |
| Cingulate cortex: area 24B | 0.52 | 0.60 | 0.65 |
| Cingulate cortex: area 32 | -0.13 | 0.89 | 0.93 |
| Dentate Gyrus of Hippocampus | -0.01 | 1.00 | 1.00 |

**Supplementary Table 4:** Whole brain volumetric analysis of mice exposed to 2 and 5 weeks of CRS as compared to vehicle.

| ROI | p Value | q Value |
|---|---|---|
| **Primary somatosensory cortex: trunk region** | **0.00** | **0.00** |
| Secondary motor cortex | 0.00 | 0.06 |
| subiculum | 0.00 | 0.06 |
| CA2Py | 0.00 | 0.09 |
| CA1Py | 0.00 | 0.14 |
| CA1Rad | 0.01 | 0.14 |
| CA2Rad | 0.01 | 0.14 |
| Lateral parietal association cortex | 0.01 | 0.14 |
| striatum | 0.00 | 0.14 |
| Medial entorhinal cortex | 0.01 | 0.22 |
| LMol | 0.02 | 0.26 |
| Cingulate cortex: area 24b | 0.02 | 0.29 |
| Cingulate cortex: area 24b' | 0.02 | 0.29 |
| Medial parietal association cortex | 0.02 | 0.30 |
| corticospinal tract/pyramids | 0.03 | 0.31 |
| medial lemniscus/medial longitudinal fasciculus | 0.03 | 0.31 |
| Frontal association cortex | 0.04 | 0.40 |
| CA20r | 0.05 | 0.40 |
| Dorsal tenia tecta | 0.05 | 0.40 |
| lobule 9 white matter | 0.04 | 0.40 |
| Primary somatosensory cortex: hindlimb region | 0.05 | 0.40 |
| trunk of simple and crus 1 white matter | 0.05 | 0.40 |
| Ventral intermediate entorhinal cortex | 0.04 | 0.40 |
| Cingulate cortex: area 32 | 0.06 | 0.41 |
| Primary somatosensory cortex: shoulder region | 0.06 | 0.41 |
| Primary somatosensory cortex: upper lip region | 0.06 | 0.41 |
| Secondary somatosensory cortex | 0.06 | 0.41 |
| lobule 9: uvula | 0.07 | 0.44 |
| lobules 4-5: culmen (ventral and dorsal) | 0.08 | 0.44 |
| mammillary bodies | 0.07 | 0.44 |
| nucleus interpositus | 0.08 | 0.44 |
| pre-para subiculum | 0.08 | 0.44 |
| thalamus | 0.07 | 0.44 |
| CA3Py Inner | 0.08 | 0.44 |
| cerebellar peduncle: inferior | 0.09 | 0.48 |
| CA3Rad | 0.11 | 0.51 |
| Dorsal intermediate entorhinal cortex | 0.11 | 0.51 |

| | | |
|---|---|---|
| GrDG | 0.11 | 0.51 |
| Olfactory bulb: mitral cell layer | 0.11 | 0.51 |
| trunk of lobules 6-8 white matter | 0.11 | 0.51 |
| medulla | 0.12 | 0.53 |
| basal forebrain | 0.14 | 0.58 |
| Primary somatosensory cortex: barrel field | 0.13 | 0.58 |
| CA10r | 0.14 | 0.60 |
| cerebellar peduncle: middle | 0.17 | 0.61 |
| lobule 3: central lobule (dorsal) | 0.17 | 0.61 |
| lobules 1-2: lingula and central lobule (ventral) | 0.16 | 0.61 |
| lobules 4-5 white matter | 0.17 | 0.61 |
| Medial preoptic nucleus | 0.17 | 0.61 |
| Olfactory bulb: internal plexiform layer | 0.17 | 0.61 |
| periaqueductal grey | 0.18 | 0.61 |
| PoDG | 0.15 | 0.61 |
| lobules 6-7 white matter | 0.19 | 0.65 |
| Primary somatosensory cortex | 0.19 | 0.65 |
| lobule 6: declive | 0.20 | 0.65 |
| habenular commissure | 0.21 | 0.67 |
| Caudomedial entorhinal cortex | 0.22 | 0.67 |
| hypothalamus | 0.22 | 0.67 |
| lobule 1-2 white matter | 0.23 | 0.67 |
| lobule 8 white matter | 0.22 | 0.67 |
| Primary motor cortex | 0.23 | 0.67 |
| trunk of arbor vita | 0.22 | 0.67 |
| Cingulate cortex: area 29c | 0.23 | 0.68 |
| SLu | 0.25 | 0.71 |
| fourth ventricle | 0.25 | 0.71 |
| colliculus: inferior | 0.27 | 0.74 |
| flocculus white matter | 0.27 | 0.74 |
| crus 2: ansiform lobule (lobule 7) | 0.29 | 0.76 |
| olfactory tubercle | 0.29 | 0.76 |
| Parietal cortex: posterior area: rostral part | 0.29 | 0.76 |
| cerebral peduncle | 0.30 | 0.76 |
| copula white matter | 0.30 | 0.76 |
| anterior commissure: pars posterior | 0.35 | 0.80 |
| CA3Py Outer | 0.34 | 0.80 |
| Cingulate cortex: area 29a | 0.36 | 0.80 |
| Frontal cortex: area 3 | 0.36 | 0.80 |
| lateral septum | 0.37 | 0.80 |

| | | |
|---|---|---|
| lobule 7: tuber (or folium) | 0.35 | 0.80 |
| lobule 8: pyramis | 0.36 | 0.80 |
| MoDG | 0.33 | 0.80 |
| Primary somatosensory cortex: dysgranular zone | 0.37 | 0.80 |
| Primary visual cortex: monocular area | 0.36 | 0.80 |
| Secondary visual cortex: mediolateral area | 0.34 | 0.80 |
| Secondary visual cortex: mediomedial area | 0.33 | 0.80 |
| Anterior olfactory nucleus | 0.39 | 0.82 |
| Cingulate cortex: area 24a' | 0.43 | 0.82 |
| Claustrum | 0.43 | 0.82 |
| cuneate nucleus | 0.45 | 0.82 |
| flocculus (FL) | 0.44 | 0.82 |
| lobule 10: nodulus | 0.42 | 0.82 |
| lobule 3 white matter | 0.47 | 0.82 |
| medial septum | 0.45 | 0.82 |
| Olfactory bulb: external plexiform layer | 0.46 | 0.82 |
| Olfactory bulb: glomerular layer | 0.44 | 0.82 |
| olfactory peduncle | 0.45 | 0.82 |
| paramedian lobule | 0.45 | 0.82 |
| Pons | 0.46 | 0.82 |
| Posterolateral cortical amygdaloid area | 0.39 | 0.82 |
| Primary auditory cortex | 0.41 | 0.82 |
| Primary somatosensory cortex: jaw region | 0.46 | 0.82 |
| Rostral amygdalopiriform area | 0.42 | 0.82 |
| subependymale zone / rhinocele | 0.47 | 0.82 |
| Ventral orbital cortex | 0.44 | 0.82 |
| crus 2 white matter | 0.47 | 0.83 |
| crus 1: ansiform lobule (lobule 6) | 0.48 | 0.84 |
| paramedian lobule (lobule 7) | 0.49 | 0.84 |
| fastigial nucleus | 0.50 | 0.85 |
| Accessory olfactory bulb: glomerular, external plexiform and mitral cell layer | 0.55 | 0.85 |
| Accessory olfactory bulb: granule cell layer | 0.54 | 0.85 |
| copula: pyramis (lobule 8) | 0.52 | 0.85 |
| Cortex-amygdala transition zones | 0.52 | 0.85 |
| dentate nucleus | 0.52 | 0.85 |
| Ectorhinal cortex | 0.54 | 0.85 |
| inferior olivary complex | 0.54 | 0.85 |
| pontine nucleus | 0.52 | 0.85 |
| Secondary auditory cortex: ventral area | 0.55 | 0.85 |

| | | |
|---|---|---|
| Temporal association area | 0.53 | 0.85 |
| trunk of lobules 1-3 white matter | 0.55 | 0.85 |
| Dorsolateral entorhinal cortex | 0.56 | 0.86 |
| anterior lobule white matter | 0.60 | 0.88 |
| cerebellar peduncle: superior | 0.60 | 0.88 |
| Claustrum: ventral part | 0.61 | 0.88 |
| Medial amygdala | 0.60 | 0.88 |
| nucleus accumbens | 0.59 | 0.88 |
| Piriform cortex | 0.58 | 0.88 |
| Secondary auditory cortex: dorsal area | 0.59 | 0.88 |
| Dorsolateral orbital cortex | 0.64 | 0.90 |
| globus pallidus | 0.64 | 0.90 |
| Primary somatosensory cortex: forelimb region | 0.64 | 0.90 |
| stria medullaris | 0.64 | 0.90 |
| simple lobule white matter | 0.66 | 0.91 |
| Amygdalopiriform transition area | 0.69 | 0.94 |
| colliculus: superior | 0.71 | 0.94 |
| crus 1 white matter | 0.71 | 0.94 |
| fasciculus retroflexus | 0.71 | 0.94 |
| Insular region: not subdivided | 0.70 | 0.94 |
| Ventral nucleus of the endopiriform claustrum | 0.70 | 0.94 |
| ventral tegmental decussation | 0.71 | 0.94 |
| cerebral aqueduct | 0.73 | 0.94 |
| internal capsule | 0.73 | 0.94 |
| Medial orbital cortex | 0.72 | 0.94 |
| CA30r | 0.73 | 0.94 |
| Dorsal nucleus of the endopiriform | 0.75 | 0.94 |
| fimbria | 0.76 | 0.94 |
| Intermediate nucleus of the endopiriform claustrum | 0.75 | 0.94 |
| Lateral orbital cortex | 0.76 | 0.94 |
| stria terminalis | 0.76 | 0.94 |
| bed nucleus of stria terminalis | 0.77 | 0.95 |
| Cingulate cortex: area 30 | 0.78 | 0.96 |
| Posteromedial cortical amygdaloid area | 0.79 | 0.96 |
| fundus of striatum | 0.81 | 0.97 |
| Primary visual cortex: binocular area | 0.81 | 0.97 |
| trunk of crus 2 and paramedian white matter | 0.81 | 0.97 |
| anterior commissure: pars anterior | 0.82 | 0.97 |
| amygdala | 0.88 | 0.97 |

| | | |
|---|---|---|
| anterior lobule (lobules 4-5) | 0.85 | 0.97 |
| Cingulate cortex: area 24a | 0.89 | 0.97 |
| Cingulate cortex: area 25 | 0.92 | 0.97 |
| Cingulate cortex: area 29b | 0.94 | 0.97 |
| Cingulum | 0.94 | 0.97 |
| Claustrum: dorsal part | 0.94 | 0.97 |
| corpus callosum | 0.93 | 0.97 |
| facial nerve (cranial nerve 7) | 0.87 | 0.97 |
| fornix | 0.91 | 0.97 |
| lateral ventricle | 0.89 | 0.97 |
| mammilothalamic tract | 0.93 | 0.97 |
| midbrain | 0.94 | 0.97 |
| Olfactory bulb: granule cell layer | 0.88 | 0.97 |
| optic tract | 0.89 | 0.97 |
| paraflocculus (PFL) | 0.90 | 0.97 |
| paraflocculus white matter | 0.86 | 0.97 |
| posterior commissure | 0.89 | 0.97 |
| Primary visual cortex | 0.90 | 0.97 |
| simple lobule (lobule 6) | 0.87 | 0.97 |
| superior olivary complex | 0.84 | 0.97 |
| third ventricle | 0.88 | 0.97 |
| Ventral tenia tecta | 0.83 | 0.97 |
| lobule 10 white matter | 0.95 | 0.97 |
| lateral olfactory tract | 0.96 | 0.98 |
| Perirhinal cortex | 0.97 | 0.98 |
| interpeduncular nucleus | 0.98 | 0.98 |
| Secondary visual cortex: lateral area | 0.99 | 1.00 |

**Supplementary Table 5:** Whole brain volumetric analysis of female mice exposed to 2 and 5 weeks of CRS as compared to vehicle.

| ROI | p Value | q Value |
|---|---|---|
| Cingulate cortex: area 29a | 0.00 | 0.36 |
| CA1Py | 0.01 | 0.64 |
| CA1Rad | 0.01 | 0.67 |
| Secondary motor cortex | 0.02 | 0.73 |
| Primary somatosensory cortex: trunk region | 0.03 | 0.73 |
| Lateral parietal association cortex | 0.03 | 0.73 |
| Primary visual cortex: monocular area | 0.05 | 0.73 |
| Secondary visual cortex: mediolateral area | 0.05 | 0.73 |
| Dorsal intermediate entorhinal cortex | 0.05 | 0.73 |
| lobule 9 white matter | 0.05 | 0.73 |
| striatum | 0.06 | 0.73 |
| medial lemniscus/medial longitudinal fasciculus | 0.06 | 0.73 |
| Medial parietal association cortex | 0.06 | 0.73 |
| Primary somatosensory cortex | 0.06 | 0.73 |
| CA3Py Outer | 0.06 | 0.73 |
| nucleus accumbens | 0.07 | 0.73 |
| Secondary visual cortex: mediomedial area | 0.08 | 0.73 |
| CA10r | 0.08 | 0.73 |
| lobule 9: uvula | 0.08 | 0.73 |
| nucleus interpositus | 0.08 | 0.73 |
| bed nucleus of stria terminalis | 0.09 | 0.73 |
| lobule 6: declive | 0.09 | 0.73 |
| Frontal cortex: area 3 | 0.09 | 0.73 |
| medulla | 0.10 | 0.73 |
| Accessory olfactory bulb: granule cell layer | 0.11 | 0.74 |
| subiculum | 0.11 | 0.74 |
| SLu | 0.12 | 0.75 |
| LMol | 0.12 | 0.75 |
| Secondary auditory cortex: dorsal area | 0.12 | 0.75 |
| Primary somatosensory cortex: hindlimb region | 0.13 | 0.75 |
| anterior commissure: pars posterior | 0.14 | 0.75 |
| Rostral amygdalopiriform area | 0.15 | 0.75 |
| Parietal cortex: posterior area: rostral part | 0.15 | 0.75 |

| | | |
|---|---|---|
| Ventral intermediate entorhinal cortex | 0.15 | 0.75 |
| corticospinal tract/pyramids | 0.15 | 0.75 |
| Primary somatosensory cortex: upper lip region | 0.15 | 0.75 |
| GrDG | 0.15 | 0.75 |
| Frontal association cortex | 0.16 | 0.78 |
| Cingulate cortex: area 30 | 0.17 | 0.81 |
| mammillary bodies | 0.19 | 0.85 |
| fastigial nucleus | 0.19 | 0.85 |
| Primary somatosensory cortex: shoulder region | 0.19 | 0.85 |
| posterior commissure | 0.20 | 0.85 |
| Lateral orbital cortex | 0.21 | 0.88 |
| Claustrum | 0.22 | 0.89 |
| PoDG | 0.24 | 0.89 |
| Cingulate cortex: area 24a | 0.24 | 0.89 |
| anterior commissure: pars anterior | 0.24 | 0.89 |
| lobules 6-7 white matter | 0.25 | 0.89 |
| CA3Rad | 0.25 | 0.89 |
| Secondary somatosensory cortex | 0.26 | 0.89 |
| Medial preoptic nucleus | 0.27 | 0.89 |
| Ventral tenia tecta | 0.27 | 0.89 |
| lobule 7: tuber (or folium) | 0.27 | 0.89 |
| CA3Py Inner | 0.28 | 0.89 |
| Primary somatosensory cortex: barrel field | 0.28 | 0.89 |
| Cingulate cortex: area 32 | 0.28 | 0.89 |
| anterior lobule white matter | 0.28 | 0.89 |
| thalamus | 0.29 | 0.89 |
| Posterolateral cortical amygdaloid area | 0.31 | 0.93 |
| Insular region: not subdivided | 0.31 | 0.93 |
| lateral olfactory tract | 0.33 | 0.93 |
| hypothalamus | 0.33 | 0.93 |
| cerebellar peduncle: superior | 0.33 | 0.93 |
| pons | 0.33 | 0.93 |
| cuneate nucleus | 0.34 | 0.93 |
| CA3Or | 0.34 | 0.93 |
| colliculus: superior | 0.36 | 0.93 |
| internal capsule | 0.36 | 0.93 |
| CA2Or | 0.37 | 0.93 |

| | | |
|---|---|---|
| CA2Rad | 0.37 | 0.93 |
| copula white matter | 0.38 | 0.93 |
| Primary somatosensory cortex: dysgranular zone | 0.39 | 0.93 |
| Medial entorhinal cortex | 0.40 | 0.93 |
| trunk of simple and crus 1 white matter | 0.40 | 0.93 |
| CA2Py | 0.40 | 0.93 |
| cerebellar peduncle: inferior | 0.40 | 0.93 |
| Olfactory bulb: mitral cell layer | 0.40 | 0.93 |
| Cingulate cortex: area 24b' | 0.40 | 0.93 |
| Cingulate cortex: area 24b | 0.43 | 0.94 |
| lobules 4-5: culmen (ventral and dorsal) | 0.44 | 0.94 |
| cerebral peduncle | 0.44 | 0.94 |
| Cingulate cortex: area 29b | 0.45 | 0.94 |
| Primary motor cortex | 0.45 | 0.94 |
| pre-para subiculum | 0.46 | 0.94 |
| simple lobule (lobule 6) | 0.46 | 0.94 |
| Olfactory bulb: glomerular layer | 0.46 | 0.94 |
| Secondary auditory cortex: ventral area | 0.46 | 0.94 |
| Primary somatosensory cortex: forelimb region | 0.47 | 0.94 |
| Anterior olfactory nucleus | 0.47 | 0.94 |
| Primary auditory cortex | 0.49 | 0.94 |
| dentate nucleus | 0.51 | 0.94 |
| mammilothalamic tract | 0.51 | 0.94 |
| basal forebrain | 0.52 | 0.94 |
| lobule 3 white matter | 0.52 | 0.94 |
| stria terminalis | 0.52 | 0.94 |
| paraflocculus (PFL) | 0.52 | 0.94 |
| trunk of arbor vita | 0.53 | 0.94 |
| olfactory tubercle | 0.54 | 0.94 |
| periaqueductal grey | 0.54 | 0.94 |
| superior olivary complex | 0.54 | 0.94 |
| MoDG | 0.55 | 0.94 |
| medial septum | 0.56 | 0.94 |
| colliculus: inferior | 0.56 | 0.94 |

| | | |
|---|---|---|
| flocculus (FL) | 0.56 | 0.94 |
| Accessory olfactory bulb: glomerular, external plexiform and mitral cell layer | 0.57 | 0.94 |
| crus 2: ansiform lobule (lobule 7) | 0.58 | 0.94 |
| lobules 4-5 white matter | 0.58 | 0.94 |
| Primary visual cortex: binocular area | 0.58 | 0.94 |
| Cingulate cortex: area 25 | 0.59 | 0.94 |
| Posteromedial cortical amygdaloid area | 0.59 | 0.94 |
| Dorsolateral entorhinal cortex | 0.60 | 0.94 |
| Secondary visual cortex: lateral area | 0.60 | 0.94 |
| Ventral orbital cortex | 0.61 | 0.94 |
| anterior lobule (lobules 4-5) | 0.61 | 0.94 |
| Olfactory bulb: internal plexiform layer | 0.62 | 0.94 |
| habenular commissure | 0.62 | 0.94 |
| Cortex-amygdala transition zones | 0.63 | 0.94 |
| lobule 1-2 white matter | 0.63 | 0.94 |
| inferior olivary complex | 0.63 | 0.94 |
| trunk of lobules 1-3 white matter | 0.63 | 0.94 |
| lobules 1-2: lingula and central lobule (ventral) | 0.65 | 0.94 |
| fourth ventricle | 0.65 | 0.94 |
| facial nerve (cranial nerve 7) | 0.66 | 0.94 |
| paraflocculus white matter | 0.66 | 0.94 |
| fasciculus retroflexus | 0.66 | 0.94 |
| Piriform cortex | 0.67 | 0.94 |
| Claustrum: ventral part | 0.67 | 0.94 |
| simple lobule white matter | 0.67 | 0.94 |
| flocculus white matter | 0.68 | 0.94 |
| Dorsal nucleus of the endopiriform | 0.69 | 0.94 |
| cerebellar peduncle: middle | 0.69 | 0.94 |
| Dorsal tenia tecta | 0.69 | 0.94 |
| Olfactory bulb: granule cell layer | 0.69 | 0.94 |
| trunk of crus 2 and paramedian white matter | 0.69 | 0.94 |
| midbrain | 0.70 | 0.94 |
| copula: pyramis (lobule 8) | 0.72 | 0.95 |

| | | |
|---|---|---|
| corpus callosum | 0.74 | 0.97 |
| Caudomedial entorhinal cortex | 0.75 | 0.97 |
| ventral tegmental decussation | 0.75 | 0.97 |
| paramedian lobule (lobule 7) | 0.76 | 0.97 |
| third ventricle | 0.76 | 0.97 |
| Ectorhinal cortex | 0.76 | 0.97 |
| Claustrum: dorsal part | 0.77 | 0.97 |
| subependymale zone / rhinocele | 0.79 | 0.98 |
| Temporal association area | 0.79 | 0.98 |
| cerebral aqueduct | 0.80 | 0.98 |
| fornix | 0.81 | 0.98 |
| Medial orbital cortex | 0.83 | 0.98 |
| fimbria | 0.83 | 0.98 |
| Intermediate nucleus of the endopiriform claustrum | 0.83 | 0.98 |
| amygdala | 0.84 | 0.98 |
| lateral septum | 0.84 | 0.98 |
| Amygdalopiriform transition area | 0.84 | 0.98 |
| Ventral nucleus of the endopiriform claustrum | 0.84 | 0.98 |
| Primary visual cortex | 0.85 | 0.98 |
| lobule 8 white matter | 0.85 | 0.98 |
| Perirhinal cortex | 0.85 | 0.98 |
| crus 2 white matter | 0.86 | 0.99 |
| fundus of striatum | 0.88 | 0.99 |
| Olfactory bulb: external plexiform layer | 0.88 | 0.99 |
| pontine nucleus | 0.89 | 0.99 |
| interpeduncIar nucleus | 0.91 | 0.99 |
| Cingulum | 0.91 | 0.99 |
| globus pallidus | 0.91 | 0.99 |
| optic tract | 0.91 | 0.99 |
| stria medullaris | 0.91 | 0.99 |
| lateral ventricle | 0.93 | 0.99 |
| trunk of lobules 6-8 white matter | 0.93 | 0.99 |
| olfactory peduncle | 0.94 | 0.99 |
| Cingulate cortex: area 24a' | 0.95 | 0.99 |
| Primary somatosensory cortex: jaw region | 0.95 | 0.99 |

| | | |
|---|---|---|
| Medial amygdala | 0.95 | 0.99 |
| crus 1 white matter | 0.96 | 0.99 |
| lobule 8: pyramis | 0.96 | 0.99 |
| Dorsolateral orbital cortex | 0.97 | 0.99 |
| paramedian lobule | 0.97 | 0.99 |
| lobule 10: nodulus | 0.97 | 0.99 |
| lobule 3: central lobule (dorsal) | 0.97 | 0.99 |
| Cingulate cortex: area 29c | 0.98 | 0.99 |
| crus 1: ansiform lobule (lobule 6) | 0.98 | 0.99 |
| lobule 10 white matter | 0.99 | 0.99 |

**Supplementary Table 6:** List of anterior cingulate cortex (ACC) direct covariance neighboring nodes in no stress, 2 weeks CRS and 5 weeks CRS groups at a density threshold of 0.13

**Regions (n=33) that covary with the ACC in the no stress group**

| No. | Name | r value |
|---|---|---|
| 6 | Anterior commissure pars posterior | 0.622 |
| 11 | Bed nucleus of stria terminalis | 0.572 |
| 22 | Cingulate cortex area 29c | 0.482 |
| 24 | Cingulate cortex area 32 | 0.742 |
| 26 | Claustrum | 0.422 |
| 27 | Claustrum dorsal part | 0.473 |
| 31 | Copula white matter | 0.622 |
| 33 | Corpus callosum | 0.641 |
| 34 | Cortex amygdala transition zones | 0.565 |
| 46 | Dorsolateral orbital cortex | 0.409 |
| 49 | Fasciculus retroflexus | 0.433 |
| 51 | Fimbria | 0.562 |
| 54 | Fornix | 0.657 |
| 55 | Frontal association cortex | 0.412 |
| 56 | Frontal cortex area 3 | 0.534 |
| 58 | Globus pallidus | 0.429 |
| 62 | Insular region not subdivided | 0.429 |
| 64 | Internal capsule | 0.505 |
| 65 | Interpedunclar nucleus | 0.492 |
| 67 | Lateral orbital cortex | 0.54 |
| 76 | Lobule 7 tuber or folium | 0.706 |
| 84 | Lobules 6-7 white matter | 0.745 |
| 86 | Mammilothalamic tract | 0.543 |
| 91 | Medial parietal association cortex | 0.411 |
| 116 | Posterior commissure | 0.522 |
| 121 | Primary motor cortex | 0.751 |
| 137 | Secondary motor cortex | 0.82 |
| 138 | Secondary somatosensory cortex | 0.557 |
| 145 | Stria terminalis | 0.446 |
| 151 | Thalamus | 0.689 |
| 153 | Trunk of crus 2 and paramedian white matter | 0.468 |
| 156 | Trunk of simple and crus 1 white matter | 0.706 |

| | | |
|---|---|---|
| 159 | Ventral orbital cortex | 0.499 |

**Regions (n=26) that covary with the ACC in the 2 weeks stress group**

| No. | Name | r value |
|---|---|---|
| 16 | Cerebellar peduncle superior | 0.505 |
| 25 | Cingulum | 0.669 |
| 33 | Corpus callosum | 0.811 |
| 51 | Fimbria | 0.552 |
| 57 | Fundus of striatum | 0.446 |
| 58 | Globus pallidus | 0.663 |
| 64 | Internal capsule | 0.746 |
| 69 | Lateral septum | 0.447 |
| 71 | Lobule 10 white matter | 0.546 |
| 78 | Lobule 8 pyramis | 0.572 |
| 79 | Lobule 9 white matter | 0.486 |
| 90 | Medial orbital cortex | 0.547 |
| 96 | Nucleus accumbens | 0.615 |
| 98 | Olfactory bulb external plexiform layer | 0.535 |
| 102 | Olfactory bulb mitral cell layer | 0.434 |
| 116 | Posterior commissure | 0.522 |
| 117 | Posterolateral cortical amygdaloid area | 0.459 |
| 122 | Primary somatosensory cortex | 0.497 |
| 124 | Primary somatosensory cortex dysgranular zone | 0.596 |
| 125 | Primary somatosensory cortex forelimb region | 0.488 |
| 126 | Primary somatosensory cortex hindlimb region | 0.486 |
| 127 | Primary somatosensory cortex jaw region | 0.484 |
| 145 | Stria terminalis | 0.696 |
| 147 | Subependymale zone rhinocele | 0.684 |
| 152 | Trunk of arbor vita | 0.504 |
| 159 | Ventral orbital cortex | 0.702 |

**Regions (n=9) that covary with the ACC in the 5 weeks stress group**

| No. | Name | r value |
|---|---|---|
| 24 | Cingulate cortex area 32 | 0.429 |
| 27 | Claustrum dorsal part | 0.462 |
| 51 | Fimbria | 0.399 |
| 56 | Frontal cortex area 3 | 0.528 |
| 62 | Insular region not subdivided | 0.562 |
| 65 | Interpeduncular nucleus | 0.527 |

| | | |
|---|---|---|
| 91 | Medial parietal association cortex | 0.462 |
| 137 | Secondary motor cortex | 0.732 |
| 141 | Secondary visual cortex mediomedial area | 0.475 |

**Supplementary Table 7:** List of amygdala direct covariance neighboring nodes in no stress, 2 weeks CRS and 5 weeks CRS groups at a density threshold of 0.08

**Regions (n=6) that covary with the amygdala in the no stress group**

| No. | Name | r value |
|---|---|---|
| 10 | Basal forebrain | 0.763 |
| 42 | Dorsal intermediate entorhinal cortex | 0.626 |
| 43 | Dorsal nucleus of the endopiriform | 0.686 |
| 87 | Medial amygdala | 0.673 |
| 113 | Piriform cortex | 0.534 |
| 134 | Rostral amygdalopiriform area | 0.564 |

**Regions (n=14) that covary with the amygdala in the 2 weeks stress group**

| No. | Name | r value |
|---|---|---|
| 9 | Anterior olfactory nucleus | 0.694 |
| 12 | Hippocampus | 0.584 |
| 13 | Caudomedial entorhinal cortex | 0.53 |
| 21 | Cingulate cortex area 29b | 0.56 |
| 29 | Colliculus inferior | 0.603 |
| 44 | Dorsal tenia tecta | 0.62 |
| 45 | Dorsolateral entorhinal cortex | 0.764 |
| 87 | Medial amygdala | 0.531 |
| 88 | Medial entorhinal cortex | 0.578 |
| 103 | Olfactory peduncle | 0.625 |
| 113 | Piriform cortex | 0.737 |
| 119 | Pre para subiculum | 0.669 |
| 150 | Temporal association area | 0.52 |
| 158 | Ventral nucleus of the endopiriform claustrum | 0.754 |

**Regions (n=16) that covary with the amygdala in the 5 weeks stress group**

| No. | Name | r value |
|---|---|---|
| 2 | Accessory olfactory bulb granule cell layer | 0.516 |
| 31 | Copula white matter | 0.603 |
| 32 | Copula pyramis lobule 8 | 0.581 |
| 34 | Cortex amygdala transition zones | 0.487 |
| 43 | Dorsal nucleus of the endopiriform | 0.61 |
| 45 | Dorsolateral entorhinal cortex | 0.715 |
| 63 | Intermediate nucleus of the endopiriform claustrum | 0.587 |
| 69 | Lateral septum | 0.643 |
| 87 | Medial amygdala | 0.686 |
| 112 | Perirhinal cortex | 0.679 |

| | | |
|---|---|---|
| 113 | Piriform cortex | 0.679 |
| 134 | Rostral amygdalopiriform area | 0.556 |
| 137 | Secondary motor cortex | 0.534 |
| 146 | Striatum | 0.519 |
| 147 | Subependymale zone rhinocele | 0.643 |
| 158 | Ventral nucleus of the endopiriform claustrum | 0.535 |
